# The individual isoforms of ORMDL, the regulatory subunit of serine palmitoyltransferase, have distinctive sensitivities to ceramide

**DOI:** 10.1101/2025.03.20.643044

**Authors:** Usha Mahawar, Deanna L Davis, Muthukumar Kannan, John Suemitsu, Christopher D Oltorik, Faheem Farooq, Raheema Fulani, Collin Weintraub, Jeremy Allegood, Binks Wattenberg

**Affiliations:** Department of Biochemistry and Molecular Biology, Virginia Commonwealth University School of Medicine, Richmond, VA 23298, USA

**Keywords:** ORMDL isoforms, Serine palmitoyltransferase (SPT), Ceramide, Sphingolipids

## Abstract

Sphingolipids play crucial roles in cell membrane structure and in multiple signaling pathways. Sphingolipid *de novo* biosynthesis is mediated by the serine palmitoyltransferase (SPT) enzyme complex. Homeostatic regulation of this complex is dependent on its regulatory subunit, the ORMDLs, of which there are three isoforms. It is well established that the ORMDLs regulate SPT activity, but it is still unclear whether the three ORMDL isoforms have distinct functions and properties. Here, we focus on understanding the physiological importance of ORMDL isoforms (ORMDL1, ORMDL2, and ORMDL3) in regulating SPT activity and sphingolipid levels. This study delves into the differential responses of the SPT complexes containing different ORMDL isoforms to cellular ceramide levels. By using the CRISPR/Cas9 gene editing tool, we have developed Hela cell lines each of which harbor only one of the three ORMDL isoforms as well as a cell line deleted for all three isoforms. Consistent with other studies, we find that deletion of all three ORMDL isoforms desensitizes SPT to ceramide and dramatically increases levels of cellular sphingolipids. In contrast, each ORMDL isoform alone is capable of regulating SPT activity and maintaining normal levels of sphingolipid. Strikingly, however, we find that each ORMDL isoform exhibits isoform-specific sensitivity to ceramide. This suggests that the inclusion of specific ORMDL isoforms into the SPT complex may accomplish a fine-tuning of sphingolipid homeostasis. The study not only emphasizes the need for further investigation into the distinct roles of ORMDL isoforms but also sheds light on their potential as therapeutic targets.

**Highlights:**

- RMDL isoforms detect varying ceramide levels to regulate SPT.
- HeLa cells, there is no compensation for the absence of the ORMDL isoform, neither at the total protein level nor at the mRNA level.

## 1. Introduction

Sphingolipids form a major and diverse class of lipids. These unique lipids are essential constituents of cell membranes and serve as signaling molecules [1]. These lipids contain a sphingoid base backbone, which is a characteristic feature of all sphingolipids [2]. This produces an amphipathic structure with a polar hydrophilic head and a non-polar hydrophobic region. The first and rate-limiting enzyme in the *de novo* sphingolipid biosynthesis pathway is serine palmitoyltransferase (SPT). This enzyme is a complex of 4 subunits, the regulatory subunits of which are known as the ORMDLs in vertebrate [3] and ORMs in yeast [4, 5]. This regulation maintains the physiological levels of sphingolipids in a cell. Cryo-electron microscopy studies have shown that the SPT/ORMDL complex in yeast [6], plants [7] and humans [8] has a ceramide binding site, which is crucial for regulating SPT activity. The binding of ceramide to the complex is specific. Only the native stereoisomer D-erythro ceramide can bind to the complex [9], and neither long-chain bases, such as sphingosine, nor higher-level sphingolipids, such as sphingomyelin, can regulate SPT. This ceramide binding is critically essential for acute and short-term SPT regulation. The ORMs/ORMDLs genes are highly conserved across different species [10]. There are three ORMDL isoforms present in a mammalian cell (ORMDL1, ORMDL2, and ORMDL3), which are highly homologous at the protein level, demonstrating over 80% sequence homology [11]. Moreover, based on siRNA silencing studies, the ORMDL isoforms appear to be functionally redundant [3]. The presence of three isoforms has raised an important question. While much has been focused on understanding how ORMDLs regulate SPT, the question remains: why do cells have three ORMDL isoforms to perform the same task? This question highlights the need for further investigation, as it could hold the key to understanding fundamental cellular processes and potential therapeutic interventions.

Disruption in ORMDL-based SPT regulation can lead to various disease conditions. For example, disruption of ORMDL-depended regulation of SPT activity leads to motor neuron defects, resulting in childhood ALS [12] and early onset of a complex form of Hereditary spastic paraplegia (HSP) [13]. Several studies have emphasized the importance of ORMDL isoforms at the mRNA level. For example, the ORMDL1 gene is upregulated in colorectal cancer [14] and in other cancer types [15]. A rare predicted loss-of-function variant of ORMDL2 is linked to age-related macular degeneration [16]. In lung cancer, ORMDL2 gene expression is significantly downregulated [17]. The ORMDL2 gene is also found to be upregulated in Immunoglobulin A nephropathy [18]. On the other hand, the most studied ORMDL isoform, ORMDL3, is found to be significantly upregulated at the RNA level in the serum samples of Alzheimer’s disease patients [19] and in patients with rheumatoid arthritis [20]. Additionally, a single nucleotide polymorphism in a non-coding region of the ORMDL3 gene has been linked to childhood asthma [21–23]. It is worth noting, however, that an alteration in mRNA expression does not necessarily correlate with protein levels. Therefore, it is crucial to verify whether changes in the mRNA level reflect changes at the protein level.

There have been limited studies on the importance of individual ORMDL isoforms. Previously, in collaboration with Dr. Spiegel and colleagues, we have shown that ORMDL triple knockout in the A549 cell line results in a significant increase in SPT activity and sphingolipid levels [24]. We also observed that knocking out ORMDL3 alone had modest effects on sphingolipid levels. Furthermore, Dr. Richard Proia and colleagues found that the whole-body ORMDL3 knockout in mice resulted in a significant increase in sphingolipid levels in brains, whereas knockout of ORMDL1 and ORMDL2 had no effect [25].

More recently, in collaboration with Dr. Xin Gong and colleagues, we showed that purified human SPT complexes containing different ORMDL isoforms demonstrate differential dose-dependent responses to ceramide [8]. Our findings indicated that ORMDL3 is more sensitive in detecting ceramide levels to regulate SPT compared to ORMDL1 and ORMDL2. The physiological findings cited above indicate isoform-specific roles of the ORMDLs. This isoform-specificity is supported by the biochemical evidence of differential responses to cellular ceramide levels.

Here, we expand on the biochemical data from purified complexes indicating differential sensitivity of SPT complexes containing different ORMDL isoforms to the more physiological setting of the SPT/ORMDL complex in the endoplasmic reticulum membrane in which this complex resides. We have used CRISPR/Cas9 gene editing to delete two of the three ORMDLs in HeLa cells and studied the effects. Our findings reveal that each ORMDL isoform senses different ceramide levels to fine-tune sphingolipid levels in a cell, which aligns with what we observed in the purified human SPT/ORMDL complex. This work represents the first step in studying the physiological importance of each individual ORMDL isoform.

## 2. Materials and Methods

### 2.1. Materials

Tissue culture: Dulbecco-modified Eagle’s medium (Gibco, # 11960-044), fetal calf serum (Gibco, #26140,079), HEPES buffer (Fisher Scientific, #BP310-1), 2mM L-glutamine (Gibco, # 25030-081), penicillin-streptomycin (Gibco, #15140-122) and Trypsin (Gibco #250-056).

CRISPR guide cloning into a lentiviral plasmid: T4 polynucleotide kinase enzyme (NEB, #M0201), T4 buffer (NEB, #B0202), nuclease-free water (Ambion #Am9938), Crispv2 Lenti plasmid (Addgene #52961), BsmB1 (NEB #R0580), DTT (NEB #7061LM), calf intestinal alkaline phosphatase (NEB #M0290), NEB buffer 3 (NEB #B6003S), GE healthcare PCR DNA and gel purification kit as per their protocol (Cytiva, #28903470), Quick ligase enzyme (# (NEB #M2200), Stellar competent bacterial cells (Takara Bio #636763).

Virus preparation: Opti-MEM (Gibco #11058021), PEI (Polysciences #239661), polyethylene glycol (Sigma #P2139-500G)

Western blotting: The following antibodies were used in this study: anti-SPTLC1 (BD Transduction Laboratories #611305), anti-SPTLC2 (Proteintech #51012-2AP), anti-SPTLC3 (ThermoFisher #PA565493), anti-ORMDL (Millipore #ABN417), anti-calnexin (Enzo ADI-SPA-860F), anti-vinculin (Proteintech #66305-1-1G), HRP-conjugated secondary antibodies for mouse (Thermo Fisher #31430) and rabbit (Thermo Fisher #31460).

Cell-based SPT activity: C-8 ceramide (Avanti polar, #860508), myriocin (Cayman Chemicals #63150), L-[^3^H]-serine (Perkin-Elmer #NET24800 or American Radiochemicals #ART0246), pyridoxal 5’phosphate (Sigma, #P9255), palmitoyl Co-A (Sigma, #P9716-10MG), L-serine (Sigma, S4500), EDTA (Thermo Fisher, #S311-500), methanol (Thermo Fisher, #A452-4), chloroform (Thermo Fisher, # C297-4), potassium hydroxide (Thermo Fisher, # P250-500), Ecolite scintillation fluid (MP Biomedicals #882475) and Beta max-scintillation fluid (MP Biomedicals #880020)

Sphingolipid analysis by Mass Spectrometry: Internal standards were purchased from Avanti Polar Lipids (Alabaster, AL).

### 2.2. Methods

#### 2.2.1. Cell culture

HeLa cells were purchased from the American Type Tissue Collection and cultured in Dulbecco-modified Eagle’s medium supplemented with 10 % fetal calf serum, HEPES buffer pH 7.), 100 U/ml penicillin-streptomycin.

#### 2.2.2. CRISPR guide RNAs designing and cloning into a lentiviral plasmid

Specific guide RNAs were designed to target exon 2 and 3 of ORMDL1, ORMDL2, and ORMDL3 isoforms (***Error! Reference source not found.***). Forward and reverse oligos for guide RNAs were synthesized from IDT, resuspended in Milli Q water for phosphorylation, and annealing by polymerase chain reaction (PCR). The reaction mixture of 10 μl was prepared, consisting of forward and reverse oligos (100 μM each), T4 polynucleotide kinase enzyme, T4 buffer, and nuclease-free water was prepared. The reaction mixture was incubated in a thermocycler (Eppendorf MasterCycler). The temperature protocol involved a 30-minute incubation at 37°C, followed by 5 minutes at 95°C, and a ramp down to 25°C at 5°C/min. Annealed oligos were diluted with nuclease-free water at 1:200 dilutions.

Crispv2 Lenti plasmid was then digested and dephosphorylated. A reaction mixture of 20 μl was prepared, which consisted of 5 µg plasmids, 1U BsmB1, 100 mM DTT, calf intestinal alkaline phosphatase, NEB buffer 3.0 and nuclease-free water. The reaction mix was incubated for 60 minutes at 37°C. The digested plasmid was run on 1% agarose gel, followed by excising out the high molecular weight band. Finally, the plasmid was eluted using a GE healthcare PCR DNA and gel purification kit as per their protocol. The diluted oligos and digested vector were ligated using Quick ligase enzyme, followed by the transformation in Stellar competent bacterial cells. The next day, at least four clones were selected, followed by a verification of successful cloning by whole plasmid sequencing.

#### 2.2.3. Virus preparation

1X 10^6^ HEK293T cells were seeded on a 100 mm tissue culture-treated dish. The next day, the cell media was replaced, and the transfection mixture was prepared. To prepare the transfection mixture, 9 μg of target plasmid and 3 μg of each viral packing plasmid (pMDLg/pRRE, pRSV-REV, and pMD2.G) were diluted with 1.5 ml of pre-warmed Opti-MEM in a 15ml falcon. In a separate 15-ml falcon, 60 μl of PEI from a 1mg/ml stock was diluted with 1.5 ml of pre-warmed Opti-MEM. Afterwards, the contents of both falcons were mixed and incubated for 20 minutes. The transfection mixture was dropwise added to the cells. The following day, virus-containing media was collected, and fresh P/S-free media was added to the cells. The above step was repeated the next day, and the virus-containing media was collected while the cells were carefully discarded.

To concentrate the virus particles, tubes containing media were centrifuged at 1200 rpm for 2 minutes at 4°C. The pellet was discarded, and collected supernatants were passed through a 0.45 µM filter. A 40% polyethylene glycol solution was prepared using 1X-PBS. 1/3 volume of the 40% PEG was added to the supernatants, maintaining the final PEG concentration at 10%. The PEG-containing supernatants were incubated for two days at a cold temperature with slow mixing. After 48 hours, the PEG-containing supernatants were centrifuged at 1500 X g for 10 minutes at 4°C. Finally, the virus-containing pellets were dissolved in 2% FBS-containing Opti-MEM. The virus particles were then stored at −80°C until needed.

#### 2.2.4. Generation of CRISPR ORMDL double and ORMDL triple knockout cell lines using HeLa cells

Hela cells were seeded at 2 × 10^6^ cells per well in a 6-well plate. The following day, the media was replaced with fresh P/S/ free media. To infect the cells with the virus, 8 µg polybrene (Sigma #TR1003-G) was diluted with 1 ml P/S free DMEM. Then, 100 µl virus particles were mixed with the polybrene-containing media and added to the cells. The plate was centrifuged at 2800 rpm for 90 minutes at 30°C. The next day, after 24 hours of transfection, the media was replaced with fresh P/S free media. The next day, the media was replaced with selection media containing one µg/ml puromycin (Sigma P4512-1ML). The cells were cultured in the selection media for at least three weeks. After three weeks, single clones were selected and cultured until they were confluent.

#### 2.2.5. Verification of CRISPR KO clones by Sanger Sequencing

To confirm successful knockouts, we extracted DNA from the single clones using a quick DNA extract solution (Bioresearch Technologies), followed by PCR amplification using specific primers (***Error! Reference source not found.***) of approximately 500 base pairs around the region where ORMDL1, ORMDL2, and ORMDL3 guide RNAs were targeted. The PCR bands were then excised and eluted from the agarose gel using GE healthcare PCR DNA and gel purification kit and sent for Sanger Sequencing (Eurofins). The sequencing results were analyzed using the CRISP-ID tool (http://crispid.gbiomed.kuleuven.be/).

#### 2.2.6. siRNA transfections

HeLa CRISPR double KO cell lines were plated at 2.5 × 10^5^ per well in a 6-well plate in Penicillin/streptomycin-free cell culture media. The day after, cells were transfected with either scrambled siRNA or siRNA against ORMDL1, ORMDL2, and ORMDL3 (***Error! Reference source not found.***) (25 pmol per well) using Lipofectamine RNAi Max (Invitrogen #13778075) as per the manufacturer’s protocol. The next day, the transfection media was replaced with P/S-free cell media. After 48 hours of transfection, cells were harvested to prepare total cell lysates.

#### 2.2.7. RNA isolation and quantitative real-time PCR

Cells were plated at 5 × 10^5^ per well in a 6-well plate. The next day, cells were harvested, and total RNA was extracted using a Trizol reagent, as previously described[26]. RNA quantity was measured using a Nano-drop, and cDNA was prepared using a high-capacity cDNA reverse transcriptase kit (Applied Biosystems #4368814) as per the manufacturer’s protocol. 7.5 ng of cDNA was used, along with pre-designed IDT-TaqMan primer probes for ORMDL1, ORMDL2, ORMDL3, SPTLC1, SPTLC2, SPTLC3, SPTssa, and SPTssb, (**Error! Reference source not found.**) as well as TaqMan Universal PCR master mix (IDT #1055772) to quantify mRNA levels. The CFX Connect real-time PCR detection system (Bio-Rad) was used to amplify the cDNA. Gene expression was calculated by the ΔΔCt method[27].

#### 2.2.8. Preparation of HeLa total cell lysates

Cells were collected after trypsinization and washed with room-temperature 1X-PBS. Then, the cell pellets were resuspended in cell lysis buffer containing 25 mM Tris, 250 mM sucrose, 1% Triton-X100, and EDTA-free protease inhibitor (Thermo Scientific #PIA)39255. The resuspended cells were passed through a 26-gauge needle 10-12 times to ensure complete cell lysis. Any remaining cell debris and unbroken cells were removed through centrifugation at 700 rpm for 5 minutes at 4°C. The Supernatants (total lysates) were carefully collected in microfuge tubes and stored at −20°C.

#### 2.2.9. Western Blotting

The protein concentration in the total lysates was measured using Bradford Reagent (Biorad) according to the manufacturer’s protocol. To prepare samples, a 5x Laemmli sample buffer was added to protein lysates and incubated at 95°C for 5 minutes. Equal amounts of protein were electrophorized on SDS/PAGE gradient gels. The proteins were transferred onto a 0.22μM PVDF membrane (Amersham #GE10600100). After transfer, blots were incubated with blocking buffers, either with a 5% blotting-grade blocker (Biorad #1706404) or 5% fat-free BSA (Fisher Scientific #BP9704100), for 2 hours at room temperature with gentle shaking. Following this, blots were incubated with anti-SPTLC1 (1:3000), anti-SPTLC2(1:2000), anti-SPTLC3 (1:2000), anti-calnexin (1:3000), anti-ORMDL (1:3000) for overnight with continuous shaking at cold. The next day, blots were washed thrice with 1x-TBST for 10 minutes each at room temperature. Meanwhile, HRP-conjugated secondary antibodies (anti-mouse and anti-rabbit) were diluted with 2.5% non-fat milk or 3% fatty acid-free BSA at 1:10,000 dilutions. Blots were then incubated with secondary antibodies overnight with continuous shaking at 4°C. The following day, the blots were washed thrice with TBST for 10 minutes each and ultrapure water for 15 minutes. Finally, the blots were visualized using an ECL-plus (Cytiva RPN2236) reagent per the manufacturer’s instructions and developed using Biorad Chemidoc, Azure Biosystems and HyblotCL film (#E3031).

#### 2.2.10. Cell-based SPT activity measurement assay

Cells were plated at 5 × 10^4^ per well on a collage-coated 24-well plate in P/S-free media. The next day, cells were transfected with either scrambled siRNA or siRNA against ORMDL1, ORMDL2, and ORMDL3 (25 pmol per well) using Lipofectamine RNAi Max according to the manufacturer’s protocol. After 48 hours of siRNA knockdown, an in-situ SPT assay was performed. A detailed protocol for an in-situ SPT assay, has been previously described[9, 28]. In brief, the cells were treated with vehicle (methanol or DMSO), C-8 ceramide, and myriocin. Subsequently, the cells were radio-labeled with [^3^H]-Serine (PerkinElmer). After radio-labeling, cells were collected with alkaline methanol, followed by sphingolipid extraction. The amount of [^3^H]-Serine incorporated into total sphingolipids was measured by liquid scintillation counting, and the results were normalized per mg of protein.

#### 2.2.11. Co-immunoprecipitation of SPT/ORMDL complex

Cells were plated at 2x 10^6^ per p100 mm dish in Penicillin/streptomycin-free media. The Next day, cells were trypsinized, collected, and washed with ice-cold 1x-PBS. Afterward, the cell pellets were resuspended in 250μl immunoprecipitation buffer consisting of 25mM-tris, 150mM Nacl, 50% glycerol, 1mM Mgcl_2_, 1mM Cacl_2_ 1x-EDTA-free protease inhibitors, and 10% (w/v) Glyco-diosgenin-GDN detergent (Anatrace #GDN1-101). The cells were then incubated for 1 hour with rotation at 4°C. Later, cells were centrifuged at 13,000 rpm for 15 min at 4°C to remove undigested cells. 50 μl supernatant was collected as input samples, and 200 μl for further processing. 1μl SPTLC2 antibody was added to 200 μl supernatant followed by overnight incubation at 4°C. The next day, supernatants were centrifuged at 13,000 rpm for 15 min at 4°C. In the meantime, 40 μl of 1:1 Protein A Sepharose 4B (Sigma #CL4B200) beads were aliquoted into Eppendorf and centrifuged for 1 min at 13,000 rpm, and the supernatant was aspirated and discarded. Carefully, water was removed from the beads, and supernatants were added. The mixture was incubated for 1 hour with rotation at 4°C. After 1 hour, supernatants were centrifuged at 13,000 rpm for 15 min at 4°C and pellets were discarded. In the meantime, 10 μl of Protein A magnetic dyna beads (Invitrogen #10001D) were aliquoted into Eppendorf and centrifuged for 1 min at 13,000 rpm to separate the buffer. Carefully, the buffer was removed from the beads, and supernatants were added. The mixture was incubated for 1 hour with rotation at 4°C. After 1 hour, supernatants were centrifuged at 13,000 rpm for 2 min at 4°C. After centrifugation, supernatants were collected and stored at −20°C. The beads were washed three times with an immunoprecipitation buffer containing 0.1% GDN, 10 minutes each wash with rotation. After the last wash, 30μl 2X-Laemmli sample buffer was added to beads, and 5μl of 5x-Laemmli sample buffer was added to 10 μl input samples. Samples were boiled at 60°C for 30 minutes, followed by SDS-PAGE electrophoresis.

#### 2.2.12. Total cell membrane isolation

Cells were seeded on at least ten p-150 tissue-culture grade dishes. Cells were harvested by trypsinization and washed with 1 x cold PBS. Cell pellets from 2 plates were resuspended in a 3.2 ml ice-cold swelling buffer (10 mM Tris pH 7.5, 15 mM KCl, 1 mM MgCl_2_) and incubated on ice for 15 minutes. Meanwhile, 1 ml 1 M sucrose, 160 µl 25× protease inhibitor, and 14 µl 200 mM EDTA pH 7.5 were added to the ice-cold Dounce homogenizer. The cells were broken by Douncing 40-45 times. Unbroken cells were removed by low-speed centrifugation at 700 rpm for 10 min at 4°C. Supernatants were collected and centrifuged at 100,000 rpm for 30 minutes at 4°C. Afterwards, pellets were collected, resuspended in 350 µl resuspension buffer (25 mM Tris pH 7.5 and 250 mM sucrose) supplemented with 1 x protease inhibitor and passed through a 16-gauge needle, then a 19-gauge needle, then a 22-needle and finally through a 26-gauge needle. Isolated membranes were aliquoted as 50 µl aliquots snap-frozen in liquid nitrogen and stored at −80 °C.

#### 2.2.13. *In vitro*-SPT activity measurement in the isolated membranes

50 µg membranes were pre-incubated in 100 µl pre-incubation buffer (50 mM HEPES, 1 mM DTT, 2 mM EDTA, 40 µM pyridoxal 5’-phosphate) for 40 minutes on ice in 2 ml microcentrifuge tubes. For ceramide dose-dependent treatment, a 1 mM C-8 ceramide stock solution was prepared by adding 20 mM C-8 ceramide in 2% fatty-acid-free BSA in 1x PBS. 1 mM C-8 was further diluted to achieve the desired concentration. In the next step, the assay was initiated by adding 100 µl pre-warmed labelling pre-mix (2 mM L-serine, 100 µM palmitoyl-CoA, and 10 µCi of [^3^H-Serine]) either in the presence of 0.5 µM, 1 µM, 2 µM, 4 µM, 5 µM and 10 µM C-8 ceramide or in the presence of control vehicle (2% fatty-acid-free BSA in 1x PBS) and 1 µM Myriocin for 1 hour at 37 °C. The assay was terminated by adding 400 µl alkaline methanol (0.7 gm KOH/100 ml methanol) and immediate vortexing. 100 µl chloroform was added to the tubes, followed by vortexing and brief centrifugation. Lipids were extracted as previously described[9, 28]. The incorporation of ^3^H-Serine in lipids was counted for 5 minutes in a Beckman-Coulter LS6500 scintillation counter.

#### 2.2.14. IC_50_ calculations

The IC_50_ values were calculated using a nonlinear regression analysis performed by GraphPad Prism version 10.0.0 for Windows, GraphPad Software, Boston, Massachusetts USA, www.graphpad.com.

#### 2.2.15. Sphingolipid analysis by Mass Spectrometry

The Lipidomics conducted in **Figure 4A** and **Figure 4B** were performed at two different time points.

##### Extraction of Sphingolipids

Tissue homogenates or cell pellets were collected in 13 × 100 mm borosilicate tubes with a Teflon-lined cap (catalogue #60827-453, VWR, West Chester, PA). Then 2 mL of CH_3_OH was added along with the internal standard cocktail (250 pmol of each species dissolved in a final total volume of 10 μl of ethanol:methanol:water 7:2:1). Standards for sphingoid bases and sphingoid base 1-phosphates were 17-carbon chain length analogs: C17-sphingosine, (2S,3R,4E)-2-aminoheptadec-4-ene-1,3-diol (d17:1-So); C17-sphinganine, (2S,3R)-2-aminoheptadecane-1,3-diol (d17:0-Sa); C17-sphingosine 1-phosphate, heptadecasphing-4-enine-1-phosphate (d17:1-So1P); and C17-sphinganine 1-phosphate, heptadecasphinganine-1-phosphate (d17:0-Sa1P). Standards for N-acyl sphingolipids were C12-fatty acid analogs: C12-Cer, N-(dodecanoyl)-sphing-4-enine (d18:1/C12:0); C12-lactosylceramide, N-(dodecanoyl)-1-β-lactosyl-sphing-4-eine (d18:1/C12:0 LacCer); C12-sphingomyelin, N-(dodecanoyl)-sphing-4-enine-1-phosphocholine (d18:1/C12:0-SM); and C12-glucosylceramide, N-(dodecanoyl)-1-β-glucosyl-sphing-4-eine (d18:1/C12:0 GlcCer). The contents were dispersed using an ultra sonicator at room temperature for 30 s. Then, 1 mL of CHCl_3_ was added, and test tubes were recapped. This single-phase mixture was incubated at 48°C overnight. The extract was centrifuged using a tabletop centrifuge, and the supernatant was removed by a Pasteur pipette and transferred to a new tube. The extract was reduced to dryness using a Speed Vac. The dried residue was reconstituted in 0.5 ml of the starting mobile phase solvent for LC-MS/MS analysis, sonicated for ca 15 sec, then centrifuged for 5 min in a tabletop centrifuge before transfer of the clear supernatant to the autoinjector vial for analysis.

##### LC-MS/MS of sphingoid bases, sphingoid base 1-phosphates, and complex sphingolipids

For LC-MS/MS analyses, a Shimadzu Nexera LC-30 AD binary pump system coupled to a SIL-30AC autoinjector and DGU20A_5R_ degasser coupled to an AB Sciex 5500 quadrupole/linear ion trap (QTrap) (SCIEX Framingham, MA) operating in a triple quadrupole mode was used. Q1 and Q3 were set to pass molecularly distinctive precursor and product ions (or a scan across multiple m/z in Q1 or Q3), using N_2_ to collisionally induce dissociations in Q2 (which was offset from Q1 by 30-120 eV); the ion source temperature set to 500 °C.These compounds were separated by reverse phase LC using a Supelco 2.1 (i.d.) x 50 mm Ascentis Express C18 column (Sigma, St. Louis, MO) and a binary solvent system at a flow rate of 0.5 mL/min with a column oven set to 35°C. Prior to injection of the sample, the column was equilibrated for 0.5 min with a solvent mixture of 95% Moble phase A1 (CH_3_OH/H_2_O/HCOOH, 58/41/1, v/v/v, with 5 mM ammonium formate) and 5% Mobile phase B1 (CH_3_OH/HCOOH, 99/1, v/v, with 5 mM ammonium formate), and after sample injection (typically 40 μL), the A1/B1 ratio was maintained at 95/5 for 2.25 min, followed by a linear gradient to 100% B1 over 1.5 min, which was held at 100% B1 for 5.5 min, followed by a 0.5 min gradient return to 95/5 A1/B1. The column was re-equilibrated with 95:5 A1/B1 for 0.5 min before the next run.

#### 2.2.16. Statistical analysis

All the experiments were performed independently at least three times. Statistical significance was determined using an unpaired two-tailed student’s t-test to compare the two groups. *p* ≤ 0.05 was considered statistically significant.

## 3. Results

### 3.1. Generation of ORMDL double knockout and triple knockout cell lines

To address the major question of why cells have three different ORMDL isoforms, we have utilized the widely used precise genome editing tool CRISP-Cas9 (Error! Reference source not found.**A)**. Using this tool, we have generated Hela cell lines, each of which has only one of the three ORMDL isoforms by deleting the other two ORMDL isoforms. We utilized guide RNAs targeting the region between exons 2 and 3 of each ORMDL isoform. The genomic location of these guide RNA in ORMDL1, ORMDL2, and ORMDL3 is shown in **(Figure 1A)**. We have also generated a cell line from which we have deleted all three ORMDL isoforms. Non-targeting guide RNAs were used to generate control cell lines.

**Figure 1:**
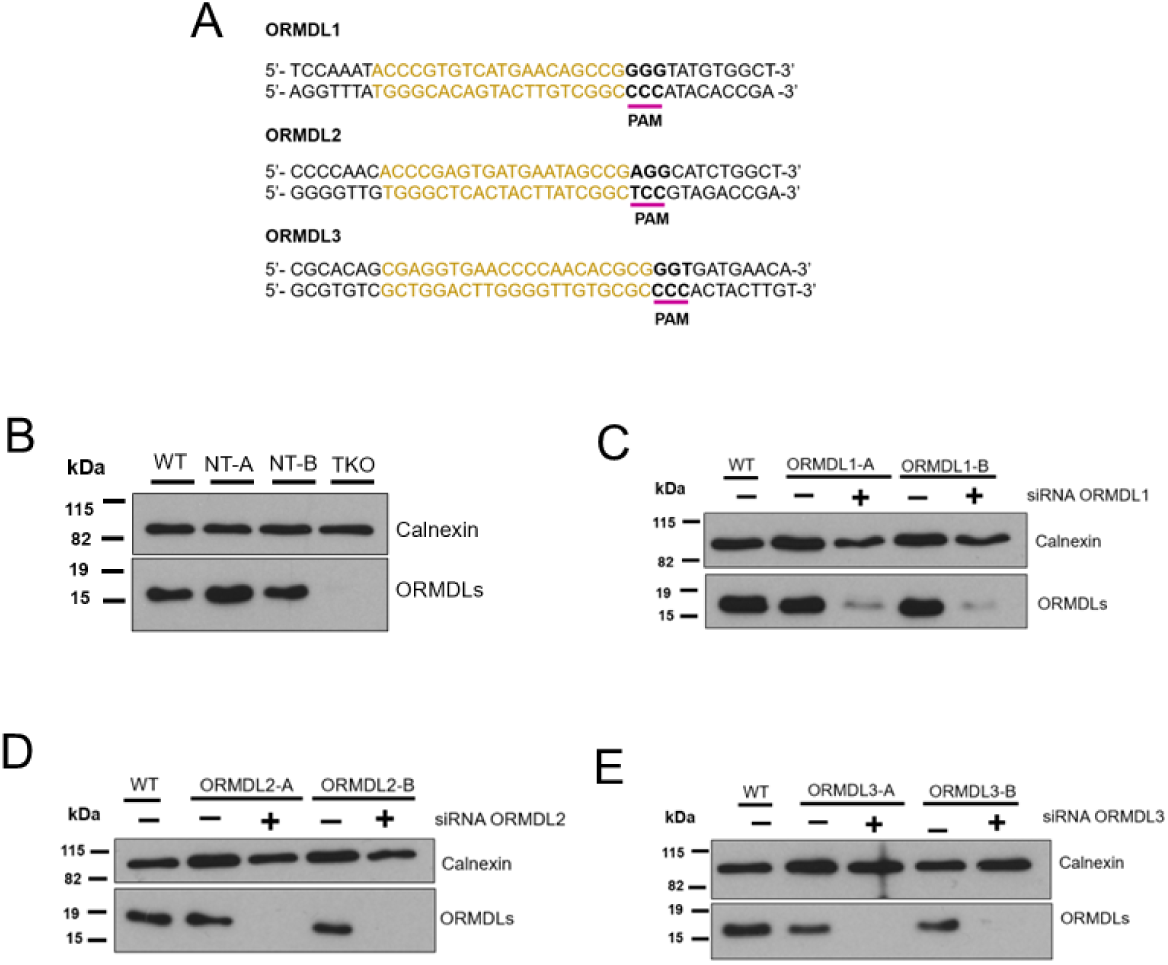
Production of ORMDL cell lines possessing only one ORMDL isoform or completely deleted ORMDLs. **A.)** Sequence of sgRNAs targeting exon 2 and 3 of ORMDL1, ORMDL2, and ORMDL3 (yellow color). The protospacer adjacent motif (PAM) (underlined in pink color) of the sgRNAs is located at the end of exon 2 and adjacent to exon 3 of ORMDL1, ORMDL2, and ORMDL3. **B.)** ORMDL protein levels in the HeLa parental cell line (WT), control cell lines (NT-A and NT-B), and ORMDL triple knockout (TKO) cell line, respectively. **C.)** Western blots to confirm the successful ORMDL2/3 double KO by siRNA knockdown of ORMDL1 in ORMDL1-A and ORMDL1-B cell lines. **D.)** Western blots to confirm the successful ORMDL1/3 double KO by siRNA knockdown of ORMDL2 in ORMDL2-A and ORMDL2-B cell lines. **E.)** Western blots to confirm the successful ORMDL1/2 double KO by siRNA knockdown of ORMDL3 in ORMDL3-A and ORMDL3-B cell lines. The western blots are representative of three individual experiments. Calnexin was used as a loading control for western blotting.

Over 40 clones were screened using the Sanger sequencing to get two positive clones for each ORMDL double-knockout cell line. To validate the deletions, approximately 500 base pairs around the targeted region of ORMDL were amplified by the PCR followed by Sanger sequencing. The sequencing reads were analyzed by the CRISPID tool (Error! Reference source not found. **and** Error! Reference source not found.) [29]. The deconvoluted sequence reads from the CRISPID were translated into protein coding region sequences using the NCBI ORFinder tool. We verified the ORMDL deletion by examining the translating the protein-coding region where guide RNAs were targeted in non-targeting control, ORMDL double and triple KO cell lines (Error! Reference source not found.**B)**. After successful verification, we isolated two positive clones for control cell lines (NT-A and NT-B), two clones of ORMDL2/3 knockout, and thus only expressing ORMDL1 (ORMDL1-A and ORMDL1-B), two clones of ORMDL1/3 knockout, and thus only expressing ORMDL2 (ORMDL2-A and ORMDL2-B), and two clones of ORMDL1/2 knockout, and thus only expressing ORMDL3 (ORMDL3-A and ORMDL3-B).

The ORMDL triple knockout (TKO) was re-verified at the protein level by western blotting **(Figure 1B)**. It should be noted that there remains a very low level of ORMDL expression in these cells **(Figure 2E)** due to the expression of a non-functional form of ORMDL1 (Error! Reference source not found.**B)**. The presence of individual ORMDL isoforms in double knockout clones at the protein level cannot be determined by western blotting because antibodies available for ORMDL cannot distinguish between different ORMDL isoforms. Therefore, to validate our isoform-specific knockout, we deleted the remaining ORMDL isoform from double knockout clones using siRNA transfection. After 48 hours of siRNA transfection, protein lysates were prepared and subjected to western blotting. By western blotting, we confirmed that the only remaining isoform in the double knockouts was the isoform not targeted by CRISPR/Cas9 **(Figure 1C-E)**.

**Figure 2:**
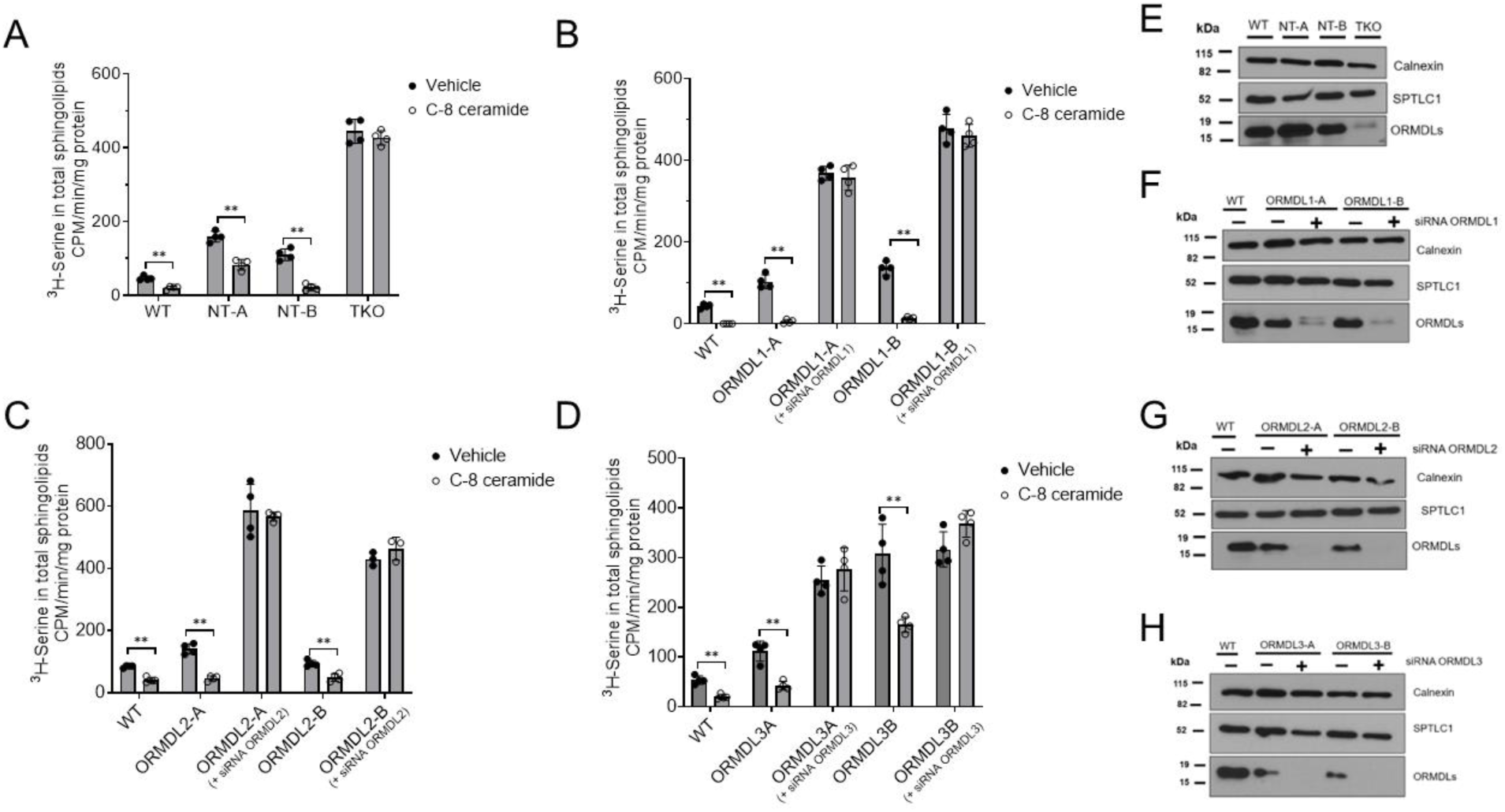
ORMDL-dependent response of SPT to exogenous ceramide is maintained when any single ORMDL isoform is expressed. SPT activity was measured in intact cells as described under “experimental procedures.” SPT activity was measured in response to a vehicle, MeOH/BSA, or 10 μM C-8 ceramide. Activity is expressed minus background, as determined with the potent SPT inhibitor myriocin. **A.)** SPT activity in WT, NT, and TKO cell lines. **B.)** SPT activity in WT, ORMDL1-A (± siRNA ORMDL1), ORMDL1-B (± siRNA ORMDL1) cell lines. **C.)** SPT activity in WT, ORMDL2-A (± siRNA ORMDL2), ORMDL2-B (± siRNA ORMDL2) cell lines. **D.)** SPT activity in WT, ORMDL3-A (± siRNA ORMDL3), ORMDL3-B (± siRNA ORMDL3) cell lines. Shown are the mean of four technical replicates, mean ± SD. Shown is one representative of three independent experiments. **E-H.)** Samples assayed for SPT activity in **A.), B.), C.), and D.)** were assessed for the presence of SPTLC1 and ORMDL protein levels by western blotting. Calnexin was used as a loading control for western blots. Statistical significance is tested by the Student’s two-tailed t-test. Asterisks denote significance p <0.01 = **.

### 3.2. Each individual ORMDL isoform can control SPT activity

The ORMDLs regulate serine palmitoyltransferase (SPT) activity by sensing cellular ceramide levels [3, 9]. Previous studies have demonstrated that deleting all three ORMDL isoforms significantly increases SPT activity [3, 24]. In the present study, SPT activity was measured by a well-characterized cell-based SPT activity measurement assay that measures the incorporation of 3H-serine into sphingolipids [28]. Responsiveness of SPT to ceramide inhibition was assessed by treating cells with C8-ceramide, a soluble form of ceramide with an 8-carbon fatty acid N-acylated to the sphingosine backbone. Basal SPT activity is dramatically increased in the ORMDL TKO cell lines compared to the wild-type and control cell lines, similar to what was previously shown in Hela Cells using siRNA depletion and in A549 cells by CRISPR/Cas9 deletion[24] **(Figure 2A)**. It was observed that the ORMDL TKO failed to inhibit SPT in the presence of C-8 ceramide, consistent with the previous report [24]. We also measured the protein level of SPTLC1, a sub-unit of SPT, by western blotting. No changes were observed in SPTLC1 protein levels in TKO compared to WT and control cell lines **(Figure 2B)**. This confirms that the increase in SPT activity is due to the release of SPT inhibition by the ORMDLs, as previously reported[9, 24]. Notably, we found that the control cell lines, NT-A and NT-B, have moderately increased SPT activity as compared to wild-type, perhaps due to clonal variation **(Figure 2A**) but respond well to ceramide inhibition. Western blotting confirmed that there are no significant changes in total protein levels of SPTLC1 and ORMDLs **(Figure 2E)**.

Next, we measured SPT activity in the absence and presence of C-8 ceramide in the ORMDL double-knockout cell lines to assess the capability of individual isoforms to regulate SPT activity. Both clones of ORMDL1 were found to regulate SPT similarly to wild-type cells. When ORMDL1 was depleted from ORMDL1-A and ORMDL1-B cell lines by siRNA knockdown, SPT activity dramatically increased, identical to the TKO cells. As expected, in ORMDL1-depleted conditions, C-8 ceramide failed to regulate SPT **(Figure 2B)**. The depletion of ORMDL1 was confirmed by western blotting **(Figure 2F)**. We also measured the SPT activity in both the clones of ORMDL2 **(Figure 2C)** and both the clones of ORMDL3 **(Figure 2D)**. Similar to ORMDL1 results, both the clones of ORMDL2 and ORMDL3 were able to regulate SPT activity similar to the wild-type cells. When the remaining ORMDL isoform was depleted from the ORMDL1/3 and ORMDL1/2 double knockout cells using siRNA knockdown, SPT activity increased, resembling that of TKO cells. ORMDL-depleted cells also failed to regulate SPT in the presence of C-8 ceramide. ORMDL depletion was also confirmed by western blotting in ORMDL2-A and ORMDL2-B cell lines **(Figure 2G)** and ORMDL3-A and ORMDL3-B cell lines **(Figure 2H).** It is worth noting that both ORMDL3 cell lines have an increased SPT activity compared to other clones, in particular ORMDL3-B. This may reflect clonal variability. Taken together, these results demonstrate that all three ORMDL isoforms can regulate SPT activity in the absence of the other two isoforms.

### 3.3. Deletion of all three ORMDL isoforms results in a significant increase in sphingolipid levels

In previous work with Dr Spiegel and colleagues, we demonstrated that ORMDL3 TKO in A549 cells leads to a significant upregulation of ceramides, sphingomyelin, and monohexosylceramides [24]. Here also, we measured the levels of C14:0, C16:0, C18:1, C18:0, C20:0, C22:0, C:24:1, C24:0, C26:1, and C:26 species of ceramide, sphingomyelin, monohexosylceramides and lactosylceramides in wildtype, control, and TKO cell lines. To calculate the total sphingolipid levels, we took the combined levels of all species of ceramide, sphingomyelin, monohexosylceramides, and lactosylceramides, which we collectively referred to as total ceramides, total sphingomyelin, total monohexosylceramides, and total lactosylceramides, respectively. The term “total monohexosylceramides” refers to a mixed population of glucosyl and galactosyl ceramides.

In control cell lines NT-A and NT-B compared to wild-type cells, we observed a moderate but statistically significant increase in total ceramides **(Figure 3A)**, total monohexosylceramides **(Figure 3D)**, total sphingomyelins **(Figure 3G)** and total lactosylceramides (Error! Reference source not found.**A)**. This may be due to increased basal SPT activity, as illustrated in Figure 2.

**Figure 3:**
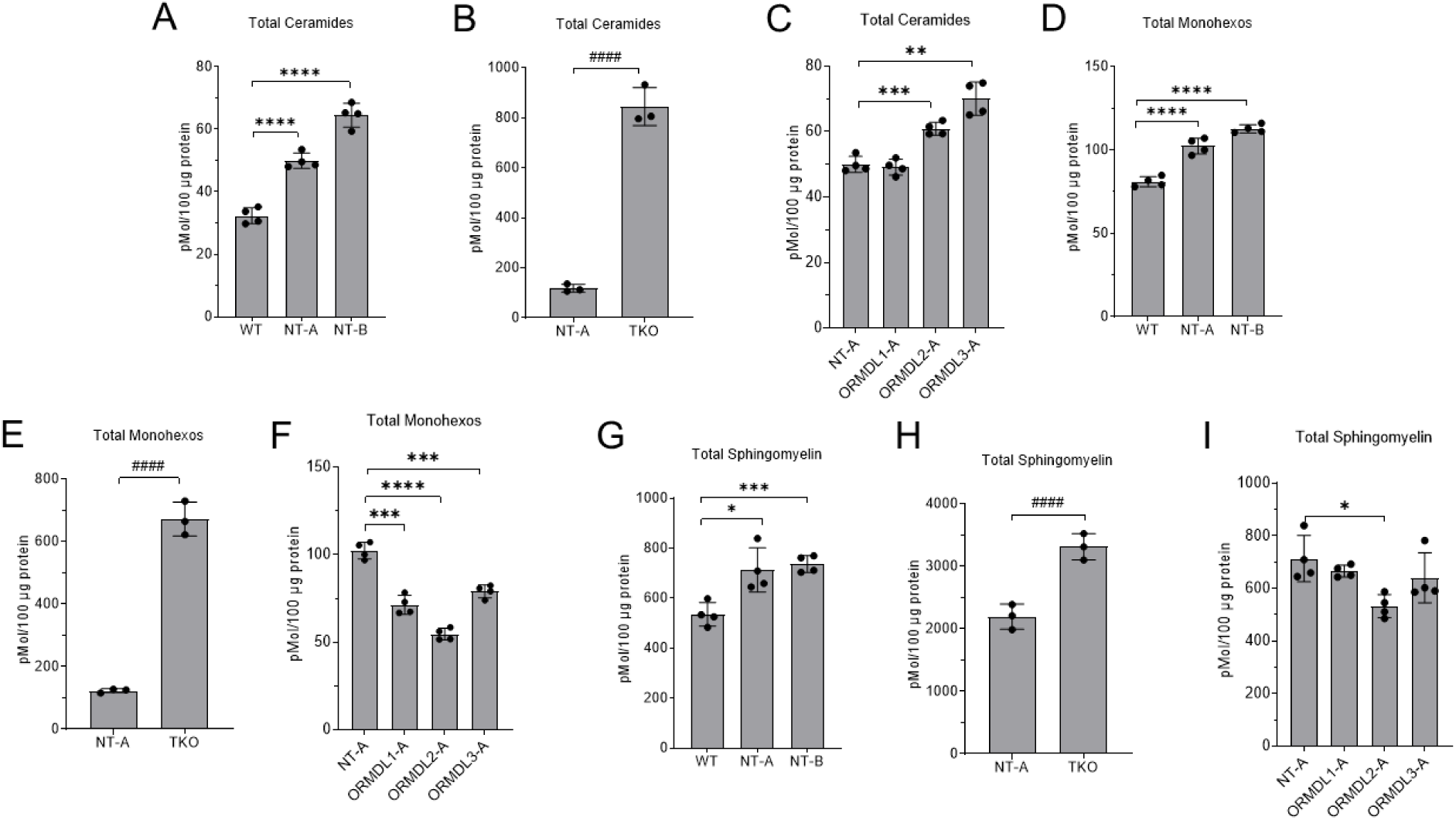
Complex sphingolipid levels are elevated in the ORMDL TKO but not ORMDL double KO cell lines. The steady-state levels of **A.), B.)** and **C.)** total ceramides**, D.), E.)** and **F.)** total monohexosylceramides, and **G.), H.)** and **I.)** total sphingomyelins in WT, NT-A, NT-B, ORMDL TKO, ORMDL1A, ORMDL2A and ORMDL3A cell lines were quantified using LC-MS/MS analysis. Data are graphed as picomoles of lipid per 100 μg of total protein. Data has been normalized to 100 μg of protein. Data is presented as mean ± SD, n=4 per cell line. Statistical significance was tested by the Student’s two-tailed t-test. Asterisks denote significance p <0.01 = *, p <0.001 = **, p <0.00001 = **** and p <0.000001 = ####. WT: wild type, NT: non-targeting, TKO: triple knockout, Monohexos: monohexosylceramides.

**Figure 4:**
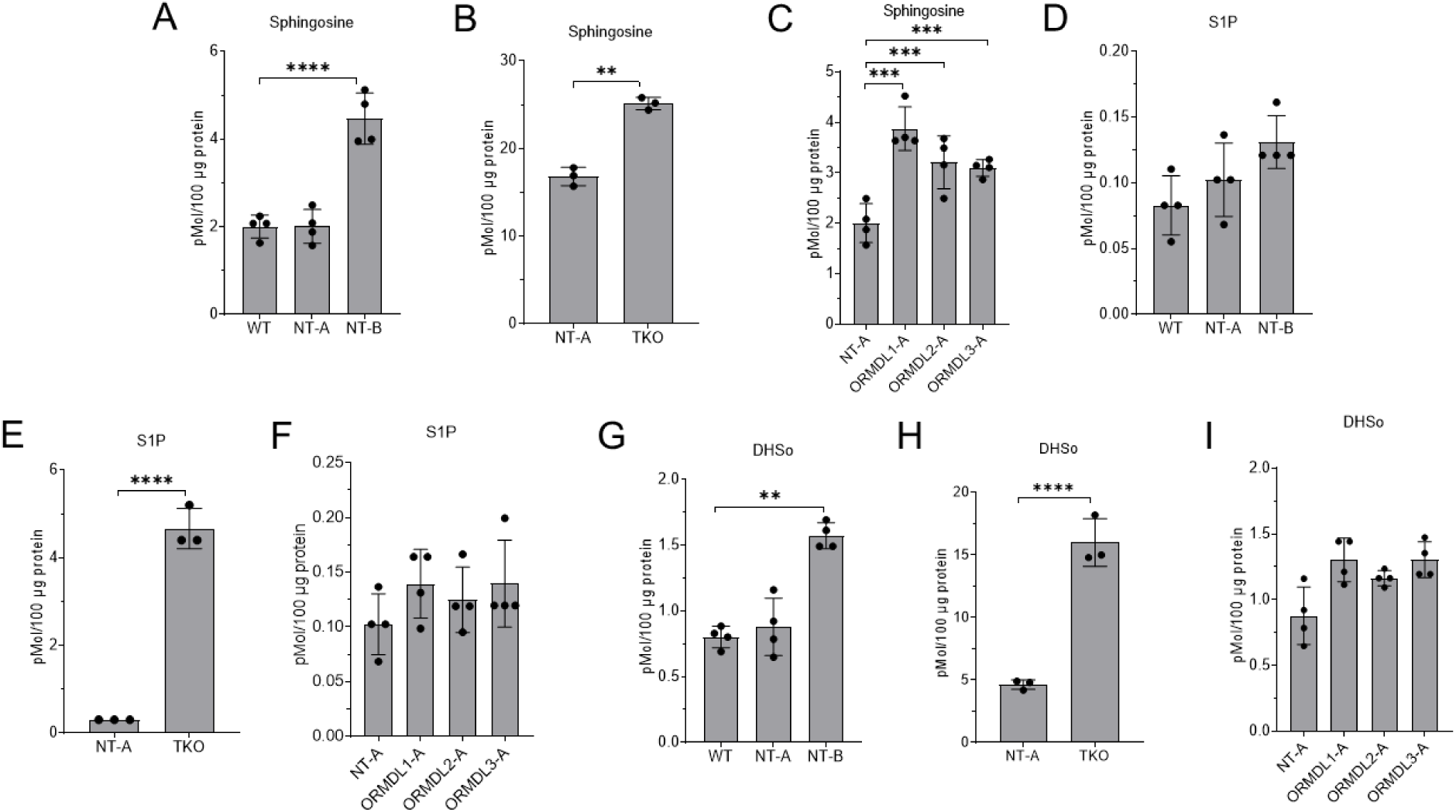
Sphingoid long-chain bases are elevated in the ORMDL TKO but not ORMDL double KO cell lines. The steady-state levels of A.), B.) and C.) sphingosine, D.), E.) and F.) sphingosine-1-phosphate, and G.), H.) and I.) dihydrosphingosine in WT, NT-A, NT-B, ORMDL TKO, ORMDL1A, ORMDL2A and ORMDL3A cell lines were quantified using LC-MS/MS analysis. Data are graphed as picomoles of lipid per 100 μg of total protein. Data is presented as mean ± SD, n=4 per cell line. Statistical significance was tested by the Student’s two-tailed t-test. Asterisks denote significance p <0.001 = **, significance p <0.0001 = ***, p <0.00001 = ***. WT: wild type, NT: non-targeting, TKO: triple knockout, S1P: sphingosine-1-phosphate, DHSo: dihydrosphingosine, and DHS1P: dihydrosphingosine-1-phosphate.

The deletion of all three ORMDLs in the TKO cell line led to an increase in total ceramides level by ∼6-fold **(Figure 3B)**. Despite dramatically increased ceramide levels, which are shown to have pro-apoptotic effects [6], TKO cells do not have any altered cell proliferation rate compared to control cells. This suggests that ORMDL TKO cells have adapted to the high cellular ceramide level conditions for their growth and proliferation. In ORMDL TKO cells, the total monohexosylceramides **(Figure 3E),** total sphingomyelins **(Figure 3H)**, and total lactosylceramides (Error! Reference source not found.**B)** also increased by ∼4.5 fold, ∼0.5 fold, and ∼1.7 fold, respectively. We also measured the levels of different acyl chain length species of ceramide, sphingomyelin, monohexosylceramides, and lactosylceramides in wild type, NT-A and NT-B control cells and TKO cell lines, respectively (Error! Reference source not found.**A-D and** Error! Reference source not found.**A-D).** In the ORMDL TKO cell lines compared to NT-A cell lines, there is a dramatic increase in all species of ceramide, monohexosylceramides, sphingomyelin, and lactosylceramides, excluding C 16:0 sphingomyelin and lactosylceramide species.

### 3.4. The steady-state levels of total ceramides, total sphingomyelins, total monohexosylceramides, and total lactosylceramides are essentially unaltered in ORMDL double KO cell lines

The SPT activity measurement assays indicate that each isoform has the potential to regulate SPT. This led us to hypothesize that each individual ORMDL isoform can control overall sphingolipid levels in the cells. In the ORMDL1-A cell line, there is no significant increase in total ceramides compared to the control cell line **(Figure 3C)**. However, in some of the double knockout cell lines, there is a very slight but statistically significant increase in total ceramides **(Figure 3C**, Error! Reference source not found.**A**). In contrast, the total monohexosylceramide levels decreased somewhat in ORMDL1-A, ORMDL2-A, and ORMDL3-A cell lines **(Figure 3F)**. In the ORMDL1-B cell line, no changes were observed in total monohexosylceramides (Error! Reference source not found.**B)**. For the ORMDL2-B cell line, the total monohexosylceramides were increased, but in ORMDL3-B cell lines, the total monohexosylceramides were decreased (Error! Reference source not found.**B)**.

Levels of total sphingomyelins and lactosylceramides remained unchanged or only slightly altered in the double knockout cell lines **(Figure 3I**, Error! Reference source not found.**C,** Error! Reference source not found.**C,** Error! Reference source not found.**D)**.

When examining the molecular species of sphingolipids in the ORMDL double KO cell lines, we did not observe any significant changes in various acyl chain length species of ceramide (Error! Reference source not found.**A and B)**, sphingomyelin (Error! Reference source not found.**A and B)**, monohexosylceramides (Error! Reference source not found.**A and B)**, and lactosylceramides (Error! Reference source not found.**A and B).** These results suggest that each ORMDL isoform can regulate overall sphingolipid levels.

### 3.5. Deletion of all three ORMDL isoforms results in a significant increase in the sphingolipid long-chain bases

Next, we measured the levels of sphingoid long-chain bases, including sphingosine (So), sphingosine-1-phosphate (S1P), and dihydrosphingosine (DHSo) in wildtype, control (NT-A and NT-B) and ORMDL TKO cell lines. Here, we observed that the NT-B cell line has a slight increase in sphingosine **(Figure 4A)** and dihydrosphingosine **(Figure 4G)** compared to the wild-type, but no changes were observed in NT-A cell lines. The levels of sphingosine-1-phosphate are not altered in both the control cell lines **(Figure 4D)**. The ORMDL TKO cell line, as expected, has a dramatic increase in sphingosine **(Figure 4B)**, sphingosine-1-phosphate **(Figure 4E),** and dihydrosphingosine **(Figure 4H)**. The levels were increased by 1.5-fold for sphingosine, 15.5-fold for sphingosine-1-phosphate, and 1.5-fold for dihydrosphingosine.

In the ORMDL double KO cell lines, sphingosine levels were significantly increased (**Figure 4C)**. In contrast, there were no consistent changes in the levels of sphingosine-1-phosphate or dihydrosphingosine in the ORMDL double-knockout cell lines (**Figure 4F**, **Figure 4I**, Error! Reference source not found.**D-F)**. These findings further validate that each ORMDL isoform can regulate the levels of sphingolipids, including intermediate metabolites of the *de novo* sphingolipid biosynthesis pathway.

### 3.6. Individual ORMDL isoforms sense different ceramide levels to regulate SPT

Our aforementioned results suggest that each ORMDL isoform can regulate SPT to maintain sphingolipid levels in the cell. We considered the possibility that although each isoform can accomplish ceramide-dependent regulation of SPT, that there might be differences in the sensitivity of each isoform to ceramide levels. Our recent publication with Gong and colleagues showed that purified human SPT/ORMDL1, SPT/ORMDL2, and SPT/ORMDL3 complexes respond to different ceramide concentrations[8]. To test this in a more physiological setting, we measured SPT activity in the presence of different ceramide concentrations in a cell-free system using membranes derived from the ORMDL knockout cell lines. The control cell lines, NT-A and NT-B, regulate SPT activity with a ceramide dose-response similar to that of wild-type cells **(Figure 5A)**. However, we observed that each ORMDL isoform had a distinct ceramide dose response for inhibiting SPT activity. ORMDL3 is more sensitive to ceramide sensing than either ORMDL1 or ORMDL2 **(Figure 5).** ORMDL2 is considerably less sensitive to ceramide sensing. We calculated the IC_50_ values to determine the concentration of C-8 ceramide in μM required to inhibit 50% SPT activity in WT, NT-A, NT-B, ORMDL1-A, ORMDL2-A, and ORMDL3-A cell lines. We observed that ORMDL3 requires 1.7 μM of C-8 ceramide to inhibit 50% SPT activity, whereas ORMDL2 needs 12 μM of C-8 ceramide to inhibit 50% SPT activity **(Figure 5C)**. Here, we confirm results from the purified complex in a physiological setting by demonstrating that in the native membrane environment, ORMDL3 is more sensitive to ceramide levels than ORMDL1 and ORMDL2, with ORMDL2 being considerably less sensitive than either ORMDL3 or ORMDL1.

**Figure 5:**
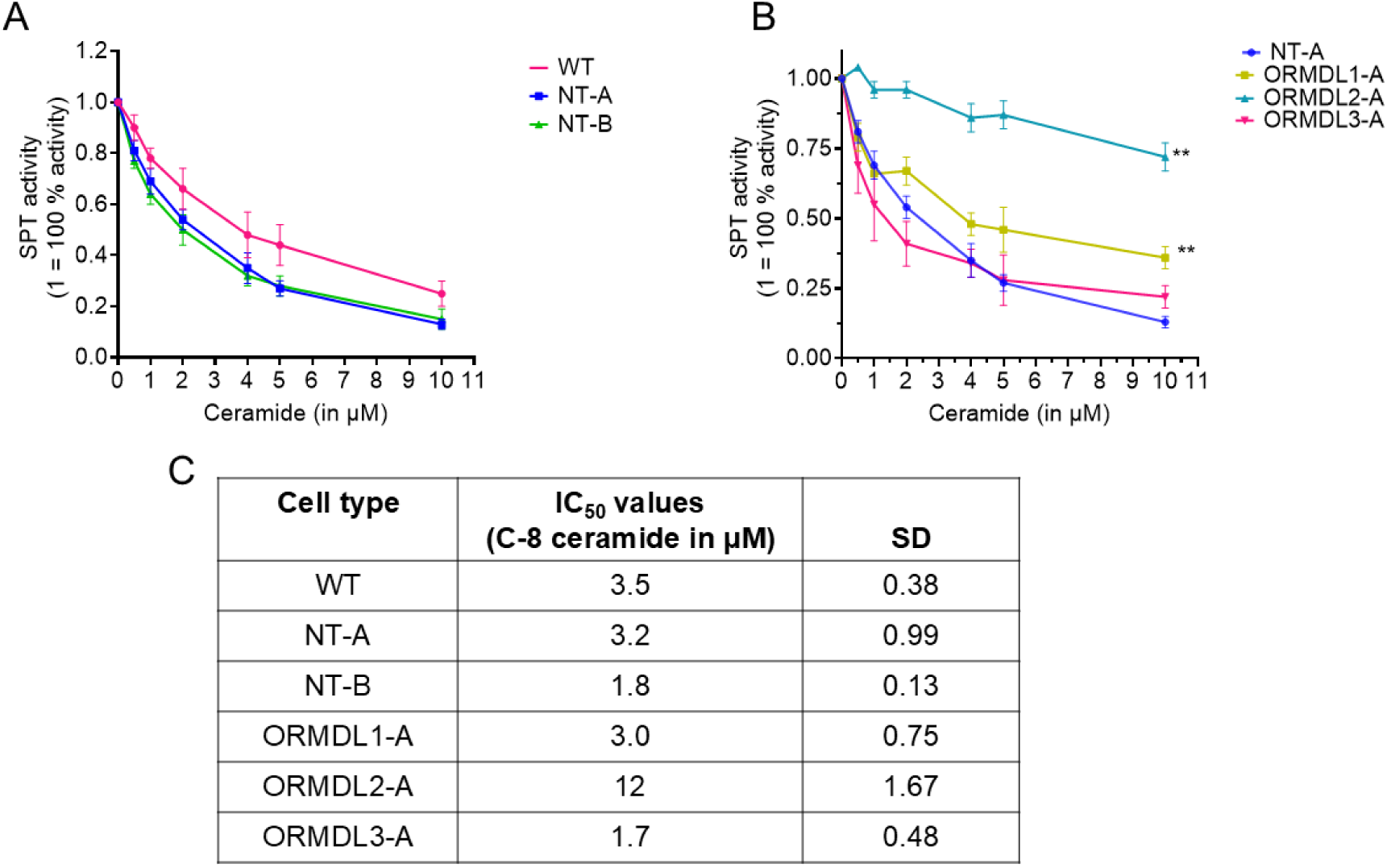
Each individual ORMDL isoform responds to different ceramide levels to regulate SPT activity. Total membranes were isolated from WT, NT-A, NT-B, ORMDL1-A, ORMDL2-A, and ORMDL3-A cell lines, and an in vitro SPT activity measurement assay was performed as described in “experimental procedures.” **A.)** In-vitro SPT activity measurements in total membranes of WT, NT-A, and NT-B cell lines. **B.)** In-vitro SPT activity measurements in total membranes of NT-A, ORMDL1-A, ORMDL2-A, and ORMDL3-A cell lines. 100 μM of total membranes were incubated with varying concentrations of BSA-conjugated C-8 ceramide (0.5 μM, 1 μM, 2 μM, 4 μM, 5 μM, and 10 μM) and tested for the ability to inhibit SPT. Untreated samples were incubated with MeOH/BSA as a control. An irreversible inhibitor of SPT, myriocin (1 μM), was used to measure the background in the assay. **C.)** IC_50_ values for NT-A ORMDL1-A, ORMDL2-A, and ORMDL3-A. Shown are the averages of quadruplicate samples, mean ± SD. Shown is the representative of at least three technical replicates. Statistical significance was tested by the Multiple T-test using GraphPad Prism Software. Asterisks denote significance p<0.001=**. Asterisks denote the significance between NT-A and ORMDL1-A, NT-A and ORMDL2-. WT: wild type, NT: non-targeting, TKO: triple knockout, KO: knockout, SPT: serine palmitoyltransferase

### 3.7. Cells do not compensate at either the protein or mRNA levels for missing ORMDL proteins

Our above results suggested that each ORMDL can regulate SPT. Next, we tested whether cells respond to the deletion of two ORMDL isoforms by upregulating the levels of the remaining isoform. We measured the total ORMDL protein levels in control cell lines, ORMDL double KO and TKO cell lines compared to wildtype cells **(Figure 6A and** Error! Reference source not found.**A)** by western blotting. The quantification of ORMDL protein levels is shown in **(Figure 6B and** Error! Reference source not found.**B).** In all cases, deletion of two of the three ORMDL isoforms results in reduced total ORMDL levels. We found that ORMDL1-A has ∼58 % and ORMDL1-B have ∼73% of ORMDL1 total protein levels, ORMDL2-A and ORMDL2-B have around ∼67% of ORMDL2 total protein levels, and ORMDL3-A and ORMDL3-B have ∼26% of ORMDL3 protein levels as compared to wildtype cells. **(Figure 6C and** Error! Reference source not found.**C)**. We also measured the total protein levels of major subunits of SPT, including SPTLC1, SPTLC2, and SPTLC3 **(Figure 6A and** Error! Reference source not found.**A)** by western blotting. The quantification of these blots showed that there are no significant changes in total protein levels in SPTLC1, SPTLC2, and SPTLC3 (Error! Reference source not found.**A-F)**. These results suggest that cells have varying total protein levels of each ORMDL isoform.

**Figure 6:**
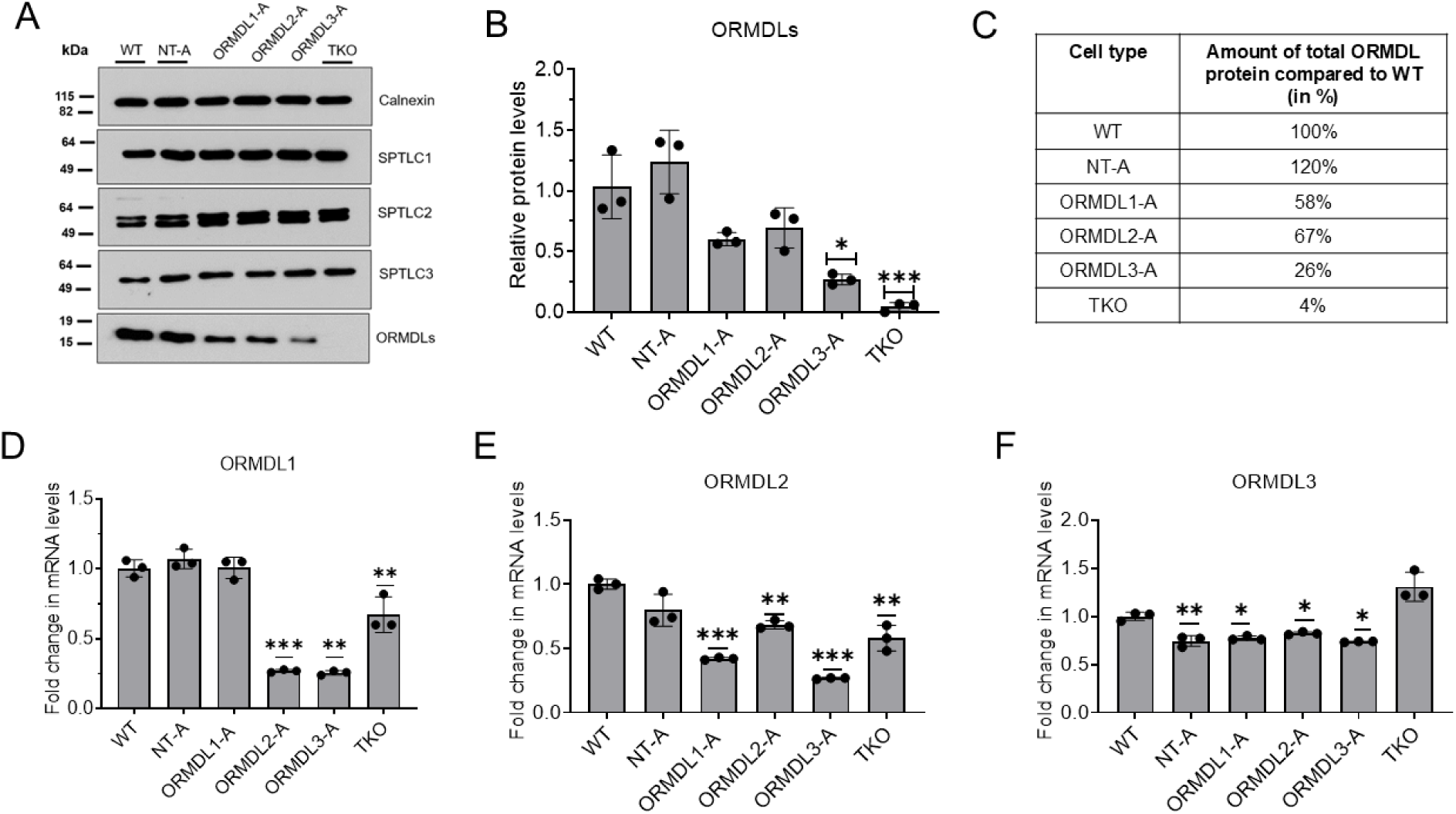
The effect of ORMDL double knockouts on total ORMDL protein and individual mRNA levels. **A.)** Representative western blot of total ORMDL, SPTLC1, SPTLC2, and SPTLC3 protein expression levels in WT, NT-A, ORMDL1-A, ORMDL2-A, ORMDL3-A, and TKO cell lines. Total cell lysates were prepared, and 10 μg of protein was loaded per well in 10-well gradient SDS-PAGE gels. Calnexin is used as a loading control. **B)** Quantification of ORMDL protein band intensity normalized to calnexin. **C.)** The total amount of OMRDL protein (in percentage) in WT, NT-A, ORMDL1-A, ORMDL2-A, ORMDL3-A, and TKO cell lines. Shown are the mean of three technical replicates, mean ± SD. **D-F.)** Total RNA was isolated from WT, NT-A, NT-B, ORMDL1-A, ORMDL1-B, ORMDL2-A, ORMDL2-B, ORMDL3-A, ORMDL3-B and TKO cell lines. RNA was reverse transcribed into cDNA, and RT-qPCR analysis was performed using 7.5 ng cDNA per reaction tube. RT-qPCR analysis of **D.)** ORMDL1, **E.)** ORMDL2, and **F.)** ORMDL3 was performed, and mRNA expression levels were normalized to hypoxanthine-guanine phosphoribosyl transferase (HPRT). The mRNA data was set relative to the control. Shown are the mean of three technical replicates, mean ± SD. Statistical significance was tested by the Student’s two-tailed t-test. Asterisks denote significance p <0.01 = *, p <0.001 = **, and p <0.0001 = ***.

To determine whether there might be compensation at the level of gene expression, we measured the mRNA expression level of ORMDL1 **(Figure 6D and** Error! Reference source not found.**D)**, ORMDL2 **(Figure 6E and** Error! Reference source not found.**E)**, and ORMDL3 **(Figure 6F and** Error! Reference source not found.**F)** in the ORMDL double KO and ORMDL triple KO cell lines. We did not observe any significant upregulation of mRNA expression levels in the ORMDL double KO and ORMDL TKO cell lines as compared to the control. Instead, we do see a decrease in the mRNA expression levels of ORMDL2 and 3 in ORMDL1 cell lines, a decrease in the mRNA expression levels of ORMDL1, 2, and 3 in ORMDL2 cell lines, and a decrease in the mRNA expression levels of ORMDL1, 2, and 3 in ORMDL3 cell lines.

We also measured the mRNA expression level of SPTLC1 (Error! Reference source not found.**A)**, SPTLC2 (Error! Reference source not found.**B)**, SPTLC3 (Error! Reference source not found.**C)**, and SPTssa (Error! Reference source not found.**D).** No significant changes were observed in the SPTLC1 and SPTLC2 mRNA expression levels. SPTLC3 mRNA levels were fluctuating up and down between both the clones of NT, ORMDL1 and ORMDL2. The total SPTLC3 protein level in the double KO cell lines suggests that the fluctuation in mRNA does not correlate with protein levels. Although there was a statistically significant decrease in SPTssa in many of the cell lines, the impact of this on SPTssa protein levels is uncertain due to the unavailability of a reliable antibody to that protein.

### 3.8. Cells maintain ORMDL levels in the SPT/ORMDL complex in the double knockout cells

As noted above, we detected reduced total ORMDL levels in the double knockout cell lines. However, there are indications that there are two ORMDL pools in cells, a pool bound to the SPT complex and a free, uncomplexed pool (addressed in more detail in Discussion). To test whether ORMDL levels are maintained in the SPT/ORMDL complex, we measured the amount of ORMDL protein bound to SPT in double KO and TKO cell lines by co-immunoprecipitation followed by western blotting **(Figure 7A and Figure 7D)**. Surprisingly, the quantification of SPT-bound ORMDL protein demonstrated that the amount of SPT-bound ORMDL in the double KO cell lines is nearly identical to levels of SPT-bound ORMDL in wildtype cells **(Figure 7G and Figure 7H)**. To verify that the SPTLC2 antibody is specifically pulling down SPTLC1 and ORMDL protein, we repeated the co-immunoprecipitation (Co-IP) experiment with control IgG. Co-IP with control IgG confirmed that our antibody is specific to SPTLC1 and ORMDL **(Figure 7C and Figure 7F)**. Very surprisingly, in TKO cell lines, we observed the presence of ORMDL protein bound to SPT. Notably, sequencing predicts an open reading frame of ORMDL3 with an insertion and multiple mutations. Based on the SPT activity measurement assay and lipidomic analysis, this is a non-functional protein. However, despite the very low level of expression of this protein, as assessed by Western blotting, it appears to be incorporated into the SPT complex. We also measured and quantified the SPT-bound SPTLC1 protein levels, and no changes were observed **(Figure 7I and Figure 7J)**. Our co-IP experiments suggest that in the ORMDL double KO cell lines, despite lower total ORMDL levels, the ORMDL content of the SPT complex is maintained.

**Figure 7:**
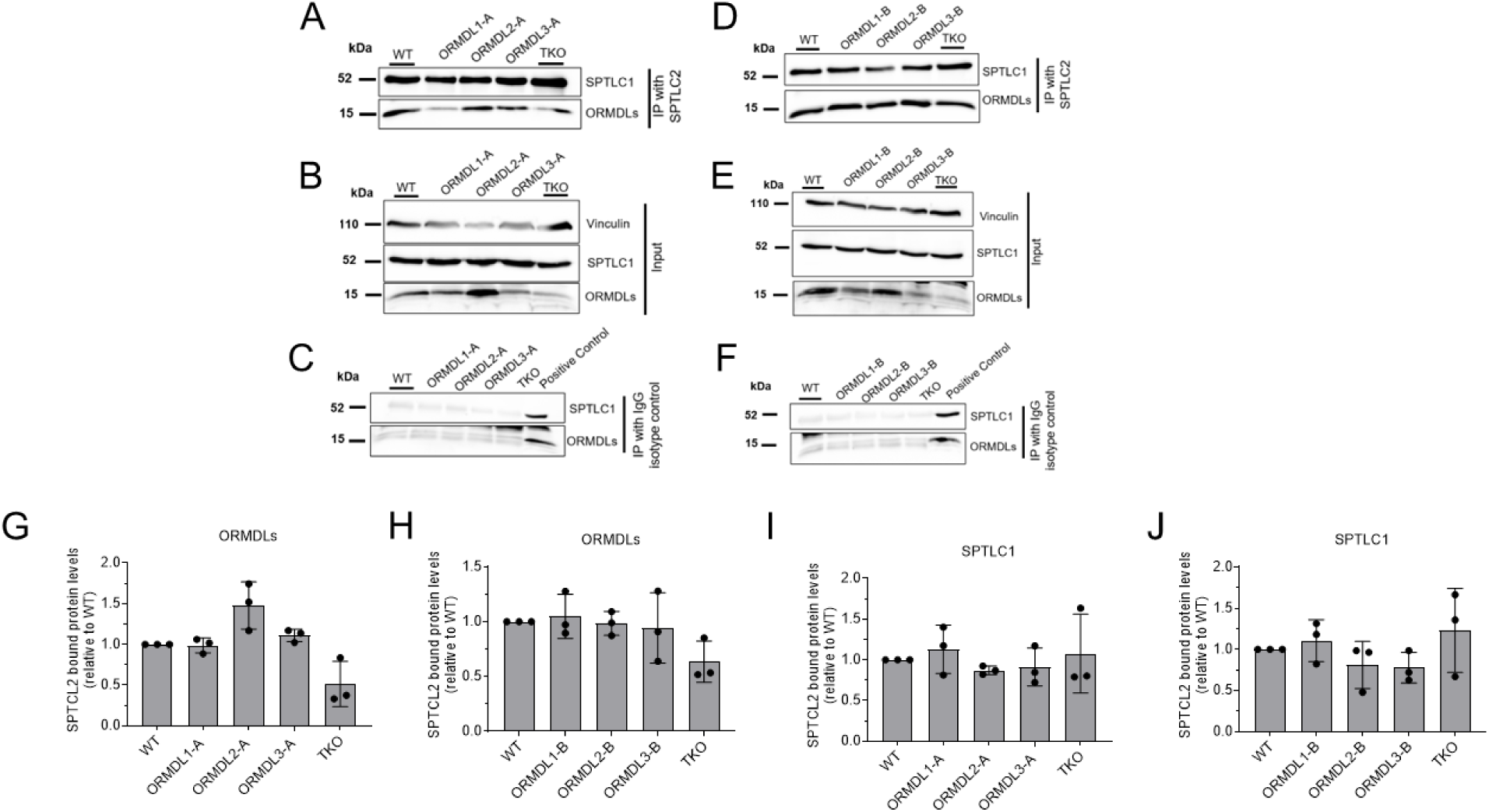
ORMDL levels are maintained in the SPT complex in ORMDL double knockout cells. **A.)** and **D.)** The SPT complex was isolated by immunoprecipitation with an antibody directed against the SPT subunit SPTLC2 as described under “experimental procedures.” Shown are the representative western blots of ORMDL and SPTLC1 protein levels in the SPT/ORMDL complex after co-immunoprecipitation. SPT complexes derived from cell lysates from **A).** WT, ORMDL1-A, ORMDL1-B, ORMDL2-A. or **D).** ORMDL2-B, ORMDL3-A, ORMDL3B, and TKO cell lines were analyzed. **B.)** and **E.)** Representative Western blots of ORMDL, SPTLC1, and Vinculin protein levels in the starting lysates (input) for the immunoprecipitations are depicted in **A)** and **D),** respectively. Vinculin was used as a loading control for the Western blot of input samples. **C.)** and **F.)** A control, non-specific IgG was used in immunoprecipitations to validate the specificity of the immunoprecipitation. Input samples are the starting material for the IP experiment. **G.)** and **H.)** Quantification of ORMDL bands from Co-IP western blot for WT, ORMDL1-A, ORMDL1-B, ORMDL2-A, ORMDL2-B, ORMDL3-A, ORMDL3B, and TKO cell lines. **I.)** and **J.)** Quantification of SPTLC1 bands from Co-IP western blot for WT, ORMDL1-A, ORMDL1-B ORMDL2-A, ORMDL2-B, ORMDL3-A, ORMDL3B, and TKO cell lines. Shown are the mean of three technical replicates, mean ± SD. Statistical significance was tested by the Student’s two-tailed t-test.

## 4. Discussion

The ORMDL proteins are crucial in controlling SPT and maintaining levels of sphingolipids within cells. Although we understand in significant detail the mechanism by which the ORMDLs regulate SPT, the individual roles of the three different ORMDL isoforms remain to be clarified. Emerging evidence from mRNA levels of the different ORMDL isoforms in tissues and different physiological conditions suggest that each isoform may play a specific role in different cell types and tissues. A major challenge in the field is understanding the significance of each ORMDL isoform at the protein level, as the commercially available ORMDL antibodies do not distinguish between the three ORMDL isoforms.

### 4.1. Differential ceramide sensitivity among different ORMDL isoforms

It is now well established that there is a ceramide binding site in the SPT/ORMDL complex, which serves to sense cellular sphingolipid levels for the homeostatic regulation of SPT activity [8]. This ceramide binding site is highly conserved among different species, including yeast and plant ORMs [6, 7, 30]. A residue in the mammalian ORMDLs (Asparagine 13) that forms an essential hydrogen bond with ceramide binding residue is highly conserved among mammalian ORMDLs, yeast ORMs, and plant ORMs [31]. Given this high level of sequence conservation, why do cells need three ORMDL isoforms? Our results revealed that each ORMDL isoform alone can efficiently regulate SPT and maintain total sphingolipid content at physiological levels. This resolves the question of whether multiple isoforms are required for the functional regulation of SPT by demonstrating that single isoforms are perfectly able to regulate SPT.

So, what explains the existence of multiple ORMDL isoforms? It is likely that cells require varying levels of the ceramide pool during different cellular events such as growth, proliferation, and differentiation. This could be accomplished by the expression of specific ORMDL isoforms with individual responses to ceramide regulation of SPT activity during these processes. For example, it has been observed that during the progression of cell division from anaphase to telophase, the levels of ceramide transfer protein (CERT) and sphingosine-1-phosphate (S1P) increase [32]. This suggests that during this transition phase, cells require increased sphingolipid synthesis. During this transition phase, the SPT/ORMDL2 complex may be well-suited for this purpose because ORMDL2 is least sensitive towards sensing ceramide, therefore allowing ceramide levels to rise before inhibition of SPT. Conversely, during non-dividing phases, sphingolipid levels must be tightly regulated, and the SPT/ORMDL3 complex is an excellent fit for this purpose.

Another possibility is that different ORMDL isoforms are present in different tissues for differential SPT regulation. For instance, sphingolipids are highly abundant lipids in the central nervous system (CNS) and skin, and dysregulation of sphingolipid metabolism leads to devastating disease conditions [33, 34]. In both these tissues, tightly controlled SPT regulation and sphingolipid metabolism are needed. ORMDL3 is well suited to the SPT complex because a slight increase in sphingolipid biosynthesis might disrupt sphingolipid metabolism and have devastating consequences for neuronal function.

### 4.2. The ORMDL content is maintained in the SPT complex despite decreased total ORMDL levels

As mentioned before, our results clearly suggest that there is no change in SPT activity or sphingolipid levels in the ORMDL double KO cells. Our findings suggest that ORMDL double KO cell lines do not compensate for missing ORMDLs at the mRNA level. These double KO cell lines have reduced total ORMDL protein levels as compared to control cells that have all three ORMDL isoforms. We find that in immunoprecipitation experiments, such as those presented here, there is a significant amount of ORMDL that does not precipitate with antibodies directed against SPTLC1 or SPTLC2. We conclude there are two pools of ORMDL; one pool represents ORMDLs incorporated into the SPT complex, and a second pool represents free or unincorporated ORMDL. We find that in the ORMDL double-knockout cell lines, despite the reduced levels of total ORMDL when compared to wild-type cells, the amount of ORMDL bound to SPT in the double KO cell lines was similar to that of wild-type cells. This indicates that the excess production of ORMDLs, resulting in a free pool of ORMDLs, ensures that the SPT complex maintains ORMDL assembly into the complex to sustain sufficient control of sphingolipid production.

## 5. Conclusion

Together, our data provide insight into the importance of each ORMDL isoform. Our results indicate that each ORMDL isoform can maintain sphingolipid homeostasis in the cells by sensing different ceramide levels. This will form the basis for a more complete understanding of the physiological role of differential expression of ORMDL isoforms in different cell types and tissues and how this impacts the crucial role that sphingolipid biosynthesis and metabolism play in the function of the organism.

## Abbreviations

SPT: Serine Palmitoyl Transferase
ORMDLs: Orosomucoid-like proteins
CRISPR: Clustered Regularly Interspaced Short Palindromic Repeats
S1P: Sphingosine-1-phosphate
HRP: Horseradish Peroxidase
PCR: Polymerase Chain Reaction

## Funding

This work was supported by NIH R21NS120128, NIH R01HL131340, the VCU Presidential Research Quest Fund, and the VCU School of Medicine Bridge Fund.

## Declaration of Competing Interest

The authors declare that they do not have any competing interests.

## Notes

### Competing Interest Statement

The authors have declared no competing interest.

## References

1. Kleuser, B., The Enigma of Sphingolipids in Health and Disease. Int J Mol Sci, 2018. 19(10).

2. Gault, C.R., L.M. Obeid, and Y.A. Hannun, An overview of sphingolipid metabolism: from synthesis to breakdown. Adv Exp Med Biol, 2010. 688: p. 1–23.

3. Siow, D.L. and B.W. Wattenberg, Mammalian ORMDL proteins mediate the feedback response in ceramide biosynthesis. J Biol Chem, 2012. 287(48): p. 40198–204.

4. Han, S., et al., Orm1 and Orm2 are conserved endoplasmic reticulum membrane proteins regulating lipid homeostasis and protein quality control. Proc Natl Acad Sci U S A, 2010. 107(13): p. 5851–6.

5. Breslow, D.K., et al., Orm family proteins mediate sphingolipid homeostasis. Nature, 2010. 463(7284): p. 1048–53.

6. Schäfer, J.H., et al., Structure of the ceramide-bound SPOTS complex. Nat Commun, 2023. 14(1): p. 6196.

7. Liu, P., et al., Mechanism of sphingolipid homeostasis revealed by structural analysis of Arabidopsis SPT-ORM1 complex. Sci Adv, 2023. 9(13): p. eadg0728.

8. Xie, T., et al., Ceramide sensing by human SPT-ORMDL complex for establishing sphingolipid homeostasis. Nat Commun, 2023. 14(1): p. 3475.

9. Davis, D.L., et al., The ORMDL/Orm-serine palmitoyltransferase (SPT) complex is directly regulated by ceramide: Reconstitution of SPT regulation in isolated membranes. J Biol Chem, 2019. 294(13): p. 5146–5156.

10. Davis, D., M. Kannan, and B. Wattenberg, Orm/ORMDL proteins: Gate guardians and master regulators. Adv Biol Regul, 2018. 70: p. 3–18.

11. Davis, D., J. Suemitsu, and B. Wattenberg, Transmembrane topology of mammalian ORMDL proteins in the endoplasmic reticulum as revealed by the substituted cysteine accessibility method (SCAM™). Biochim Biophys Acta Proteins Proteom, 2019. 1867(4): p. 382–395.

12. Mohassel, P., et al., Childhood amyotrophic lateral sclerosis caused by excess sphingolipid synthesis. Nat Med, 2021. 27(7): p. 1197–1204.

13. Srivastava, S., et al., SPTSSA variants alter sphingolipid synthesis and cause a complex hereditary spastic paraplegia. Brain, 2023. 146(4): p. 1420–1435.

14. Wang, Q., et al., ORMDL1 is upregulated and associated with favorable outcomes in colorectal cancer. Transl Oncol, 2021. 14(10): p. 101171.

15. Zhu, T., et al., Expression Patterns and Prognostic Values of ORMDL1 in Different Cancers. Biomed Res Int, 2020. 2020: p. 5178397.

16. Kwong, A., et al., Whole genome sequencing of 4,787 individuals identifies gene-based rare variants in age-related macular degeneration. Hum Mol Genet, 2024. 33(4): p. 374–385.

17. Maiuthed, A., O. Prakhongcheep, and P. Chanvorachote, Microarray-based Analysis of Genes, Transcription Factors, and Epigenetic Modifications in Lung Cancer Exposed to Nitric Oxide. Cancer Genomics Proteomics, 2020. 17(4): p. 401–415.

18. Xu, J., et al., Identification of blood-based key biomarker and immune infiltration in Immunoglobulin A nephropathy by comprehensive bioinformatics analysis and a cohort validation. J Transl Med, 2022. 20(1): p. 145.

19. Shao, Y., et al., The inhibition of ORMDL3 prevents Alzheimer’s disease through ferroptosis by PERK/ATF4/HSPA5 pathway. IET Nanobiotechnol, 2023. 17(3): p. 182–196.

20. Ye, W., et al., A Common Functional Variant at the Enhancer of the Rheumatoid Arthritis Risk Gene ORMDL3 Regulates its Expression Through Allele-Specific JunD Binding. Phenomics, 2023. 3(5): p. 485–495.

21. Ober, C., et al., Expression quantitative trait locus fine mapping of the 17q12-21 asthma locus in African American children: a genetic association and gene expression study. Lancet Respir Med, 2020. 8(5): p. 482–492.

22. Toncheva, A.A., et al., Childhood asthma is associated with mutations and gene expression differences of ORMDL genes that can interact. Allergy, 2015. 70(10): p. 1288–99.

23. Wills-Karp, M., At last - linking ORMDL3 polymorphisms, decreased sphingolipid synthesis, and asthma susceptibility. J Clin Invest, 2020. 130(2): p. 604–607.

24. Green, C.D., et al., CRISPR/Cas9 deletion of ORMDLs reveals complexity in sphingolipid metabolism. J Lipid Res, 2021. 62: p. 100082.

25. Clarke, B.A., et al., The Ormdl genes regulate the sphingolipid synthesis pathway to ensure proper myelination and neurologic function in mice. Elife, 2019. 8.

26. Davis, D.L., et al., Dynamics of sphingolipids and the serine palmitoyltransferase complex in rat oligodendrocytes during myelination. J Lipid Res, 2020. 61(4): p. 505–522.

27. Livak, K.J. and T.D. Schmittgen, Analysis of relative gene expression data using real-time quantitative PCR and the 2(-Delta Delta C(T)) Method. Methods, 2001. 25(4): p. 402–8.

28. Kannan, M., et al., Preparation of HeLa Total Membranes and Assay of Lipid-inhibition of Serine Palmitoyltransferase Activity. Bio Protoc, 2020. 10(12): p. e3656.

29. Dehairs, J., et al., CRISP-ID: decoding CRISPR mediated indels by Sanger sequencing. Sci Rep, 2016. 6: p. 28973.

30. Xie, T., et al., Collaborative regulation of yeast SPT-Orm2 complex by phosphorylation and ceramide. Cell Rep, 2024. 43(2): p. 113717.

31. Mughram, M.H.A., G.E. Kellogg, and B.W. Wattenberg, Three kingdoms and one ceramide to rule them all. A comparison of the structural basis of ceramide-dependent regulation of sphingolipid biosynthesis in animals, plants, and fungi. Adv Biol Regul, 2023: p. 101010.

32. Voelkel-Johnson, C., Sphingolipids in embryonic development, cell cycle regulation, and stemness - Implications for polyploidy in tumors. Semin Cancer Biol, 2022. 81: p. 206–219.

33. Borodzicz, S., et al., The role of epidermal sphingolipids in dermatologic diseases. Lipids Health Dis, 2016. 15: p. 13.

34. Leal, A.F., et al., Sphingolipids and their role in health and disease in the central nervous system. Adv Biol Regul, 2022. 85: p. 100900.

35. The graphical abstract figure was created in BioRender. Mahawar, U. (2025) http://BioRender.com/f68i214

