## Supplementary Figure Legends for "The individual isoforms of ORMDL, the regulatory subunit of serine palmitoyltransferase, have distinctive sensitivities to ceramide"

**Supplementary Figure 1: Verification of ORMDL TKO at the genomic level.** The genomic DNA sequences were determined around the region of the RNA guides used to generate knockouts **A.)** HeLa WT, **B.)** ORMDL TKO, **C.)** NT-A and **D.)** NT-B cell lines, respectively. ~350 base pair regions around the guide RNA target were amplified using PCR. PCR bands were eluted and sent for the Sanger sequencing. The Sanger sequencing reads were deconvoluted by CRISPID and compared to the reference sequence. The reference sequence (the top sequence in the figure) is genomic DNA sequences ± 400 base pairs around the region where guide RNAs for ORMDL1, ORMDL2, and ORMDL3 were targeted. Guide RNAs were highlighted with yellow color. The bottom sequence is the deconvoluted sequence from the original sequencing traces.

**Supplementary Figure 2: Verification of ORMDL double knockouts at the genomic level.** The genomic DNA sequences were determined around the region of the RNA guides used to generate knockouts **A.)** ORMDL1-A cell line, **B.)** ORMDL1-B, **C.)** ORMDL2-A **D.)** ORMDL2-B, **E.)** ORMDL3-A and **F.)** ORMDL3-B cell lines, respectively. ~350 base pair regions around the guide RNA target were amplified using PCR. PCR bands were eluted and sent for the Sanger sequencing. The Sanger sequencing reads were deconvoluted by CRISPID and compared to the reference sequence. The reference sequence (the top sequence in the figure) is genomic DNA sequences ± 400 base pairs around the region where guide RNAs for ORMDL1, ORMDL2, and ORMDL3 were targeted. Guide RNAs were highlighted with yellow color. The bottom sequence is the deconvoluted sequence from the original sequencing traces.

**Supplementary Figure 3: Predicted protein sequences of ORMDL double knockouts.** **A.)** Schematic representation of CRISPR Cas9 machinery. **B.)** The sequencing reads were translated into the protein-coding region to verify knockout at the protein level in the WT, control cell lines (NT-A and NT-B), ORMDL1-A and ORMDL1-B cell lines (ORMDL2/3 KO), ORMDL2-A and ORMDL2-B cell lines (ORMDL1/3 KO), and ORMDL3-A and ORMDL3-B cell line (ORMDL1/2 KO)**.** sgRNA targeting amino acids were highlighted in yellow color.

**Supplementary Figure 4: Lactosylceramides level in wildtype, control, TKO and ORMDL double knockout cell lines.** Steady-state levels of total lactosylceramides in **A.)**  WT, NT-A, and NT-B, **B.)** NT-A and TKO, **C.)** NT-A, ORMDL1-A, ORMDL2-A, and ORMDL3-A and **D.)** NT-B, ORMDL1-B, ORMDL2-B, and ORMDL3-B cell lines were quantified using LC-MS/MS analysis. Data are graphed as picomoles of lipid per 100 μg of total protein. Data is presented as mean ± SD, n=4 per cell line. Statistical significance was tested by the Student’s two-tailed t-test. Asterisks denote significance p <0.01 = *.

**Supplementary Figure 5: The steady-state levels of different acyl chain length sphingolipid species in control cell lines.** The levels of C14:0, C16:0, C18:1, C18:0, C20:0, C22:0, C:24:1, C24:0, C26:1, and C26:0 **A.)** ceramide species, **B.)** sphingomyelin species, **C.)** monohexosylceramide species, and **D.)** lactosylceramide species were quantified in WT, NT-A, NT-B cell lines using LC-MS/MS analysis. Data are graphed as picomoles of lipid per 100 μg of total protein. Data are presented as mean ± SD, n=4 per cell line.

**Supplementary Figure 6: Comparison of the steady-state levels of different acyl chain length sphingolipid species in WT vs ORMDL3 TKO cell lines.** The levels of C14:0, C16:0, C18:1, C18:0, C20:0, C22:0, C:24:1, C24:0, C26:1, and C26:0 **A.)** ceramide species, **B.)** sphingomyelin species, and **C.)** monohexosylceramides species were quantified in WT and TKO cell lines using LC-MS/MS analysis. Data are graphed as picomoles of lipid per 100 μg of total protein. Data are presented as mean ± SD, n=4 per cell line.

**Supplementary Figure 7: The complex sphingolipid and sphingoid long-chain base levels in the Clone B of ORMDL double KO cell lines.** Steady-state levels of **A.)** total ceramides, **B.)** total monohexosylceramides, **C.)** total sphingomyelin, **D.)** sphingosine, **E.)** sphingosine-1-phosphate and **F.)** dihydrosphingosine in NT-B, ORMDL1-B, ORMDL2-B, and ORMDL3-B cell lines were quantified using LC-MS/MS analysis. Data are graphed as picomoles of lipid per 100 μg of total protein. Data is presented as mean ± SD, n=4 per cell line. Statistical significance was tested by the Student’s two-tailed t-test. Asterisks denote significance p <0.001 = **, p <0.00001 = ***.

**Supplementary Figure 8: The steady-state level of different acyl chain length ceramide species in the ORMDL double knockout cell lines as compared to a control cell line.** The ceramide levels of **(A)** C14:0, C16:0, C18:1, C18:0, and C20:0, C22:0, C:24:1, C24:0, C26:1, and C26:0 was quantified in NT-A, ORMDL1-A, ORMDL2-A and ORMDL3-A using LC-MS/MS analysis. The ceramide levels of **(B)** C14:0, C16:0, C18:1, C18:0, and C20:0, C22:0, C:24:1, C24:0, C26:1, and C26:0 was quantified in NT-B, ORMDL1-B, ORMDL2-B and ORMDL3-B using LC-MS/MS analysis. Data are graphed as picomoles of lipid per 100 μg of total protein.

**Supplementary Figure 9: The steady-state level of different acyl chain length sphingomyelin species in the ORMDL double knockout cell lines as compared to a control cell line.** The sphingomyelin levels of **(A)** C14:0, C16:0, C18:1, C18:0, and C20:0, C22:0, C:24:1, C24:0, C26:1, and C26:0 was quantified in NT-A, ORMDL1-A, ORMDL2-A and ORMDL3-A using LC-MS/MS analysis. The ceramide levels of **(B)** C14:0, C16:0, C18:1, C18:0, and C20:0, C22:0, C:24:1, C24:0, C26:1, and C26:0 was quantified in NT-B, ORMDL1-B, ORMDL2-B and ORMDL3-B using LC-MS/MS analysis. Data are graphed as picomoles of lipid per 100 μg of total protein. Data have been normalized to 100 μg of phosphate. Data is presented as mean ± SD, n=4 per cell line. Statistical significance was tested by the Student’s two-tailed t-test. NT: non-targeting, KO: knockout.

**Supplementary Figure 10: The steady-state level of different acyl chain length monohexosylceramide species in the ORMDL double knockout cell lines as compared to a control cell line.** The monohexosylceramide level of **(A)** C14:0, C16:0, C18:1, C18:0, and C20:0, C22:0, C:24:1, C24:0, C26:1, and C26:0 was quantified in NT-A, ORMDL1-A, ORMDL2-A and ORMDL3-A using LC-MS/MS analysis. The ceramide levels of **(B)** C14:0, C16:0, C18:1, C18:0, and C20:0, C22:0, C:24:1, C24:0, C26:1, and C26:0 was quantified in NT-B, ORMDL1-B, ORMDL2-B and ORMDL3-B using LC-MS/MS analysis. Data are graphed as picomoles of lipid per 100 μg of total protein. Data are presented as mean ± SD, n=4 per cell line.

**Supplementary Figure 11: The steady-state level of different acyl chain length lactosylceramide species in the ORMDL double knockout cell lines as compared to a control cell line.** The lactosylceramide levels of **(A)** C14:0, C16:0, C18:1, C18:0, and C20:0, C22:0, C:24:1, C24:0, C26:1, and C26:0 was quantified in NT-A, ORMDL1-A, ORMDL2-A and ORMDL3-A using LC-MS/MS analysis. The ceramide levels of **(B)** C14:0, C16:0, C18:1, C18:0, and C20:0, C22:0, C:24:1, C24:0, C26:1, and C26:0 was quantified in NT-B, ORMDL1-B, ORMDL2-B and ORMDL3-B using LC-MS/MS analysis. Data are graphed as picomoles of lipid per 100 μg of total protein. Data is presented as mean ± SD, n=4 per cell line.

**Supplementary Figure 12: Total ORMDL, SPTLC1, SPTLC2 and SPTLC3 protein levels in the ORMDL double and triple KO cell lines.** **A.)** Representative western blot of total ORMDL, SPTLC1, SPTLC2, and SPTLC3 protein expression levels in WT, NT-B, ORMDL1-B, ORMDL2-B, ORMDL3-B, and TKO cell lines. Total cell lysates were prepared, and 10 μg of protein was loaded per well in 10-well gradient SDS-PAGE gels. Calnexin is used as a loading control. **B)** Quantification of ORMDL protein band intensity normalized to calnexin. **C.)** The total amount of OMRDL protein (in percentage) in WT, NT-B, ORMDL1-B, ORMDL2-B, ORMDL3-B, and TKO cell lines. Shown are the mean of three technical replicates, mean ± SD. RT-qPCR analysis of **D.)** ORMDL1, **E.)** ORMDL2, and **F.)** ORMDL3 was performed, and mRNA expression levels were normalized to hypoxanthine-guanine phosphoribosyl transferase (HPRT). The mRNA data was set relative to the control. Shown are the mean of three technical replicates, mean ± SD. Statistical significance was tested by the Student’s two-tailed t-test. Asterisks denote significance p <0.01 = *, p <0.001 = **, and p <0.0001 = ***.

**Supplementary Figure 13: Quantification of total SPTLC1, SPTLC2, and SPTLC3 protein levels in the ORMDL double and triple KO cell lines.** Quantification of total **A.)** SPTLC1, **B.)** SPTLC2 and **C.)** SPTLC3 protein band intensity normalized to calnexin.

**Supplementary Figure 14: The mRNA expression levels of SPTLC1, SPTLC2, SPTLC3, and SPTssa in the ORMDL double and triple KO cell lines.** Total RNA was isolated from WT, NT-A, NT-B, ORMDL1-A, ORMDL1-B, ORMDL2-A, ORMDL2-B, ORMDL3-A, ORMDL3-B and TKO cell lines. RNA was reverse transcribed into cDNA, and RT-qPCR analysis was performed using 7.5 ng cDNA per reaction tube. RT-qPCR analysis of **A.)** SPTLC1, **B.)** SPTLC2, **C.)** SPTLC3 and **D.)** SPTssa was performed, and mRNA expression levels were normalized to HPRT. The mRNA data was set relative to the control. Shown are the mean of three technical replicates, mean ± SD. Statistical significance for a difference from WT was tested by the Student’s two-tailed t-test. Asterisks denote significance p <0.01 = *, p <0.001 = **, p <0.0001 = *** and p <0.00001 = ****.
