## Supplementary Figures and Tables for "The individual isoforms of ORMDL, the regulatory subunit of serine palmitoyltransferase, have distinctive sensitivities to ceramide"

### Slide 1
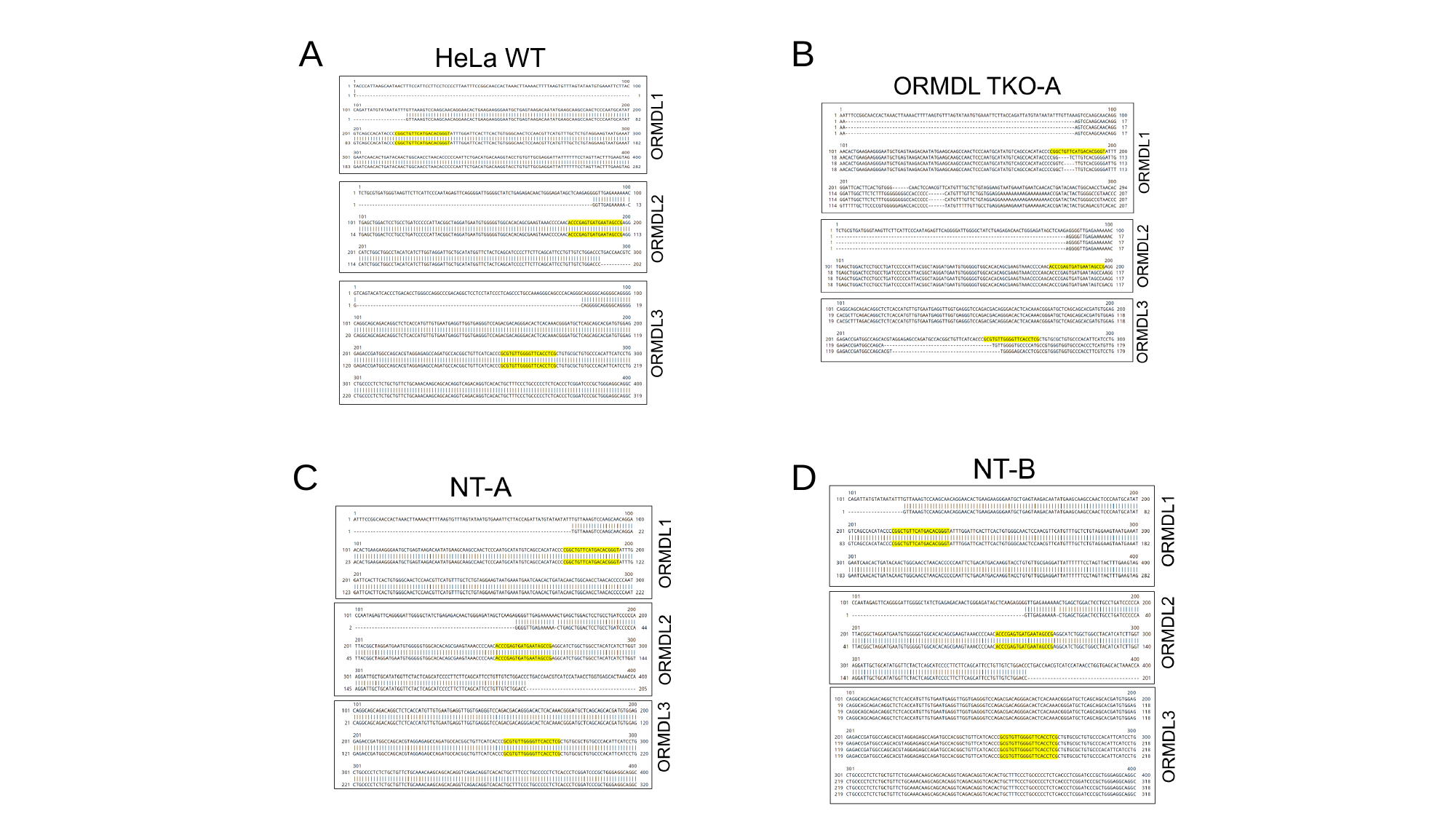

A
B
D
C

### Slide 2
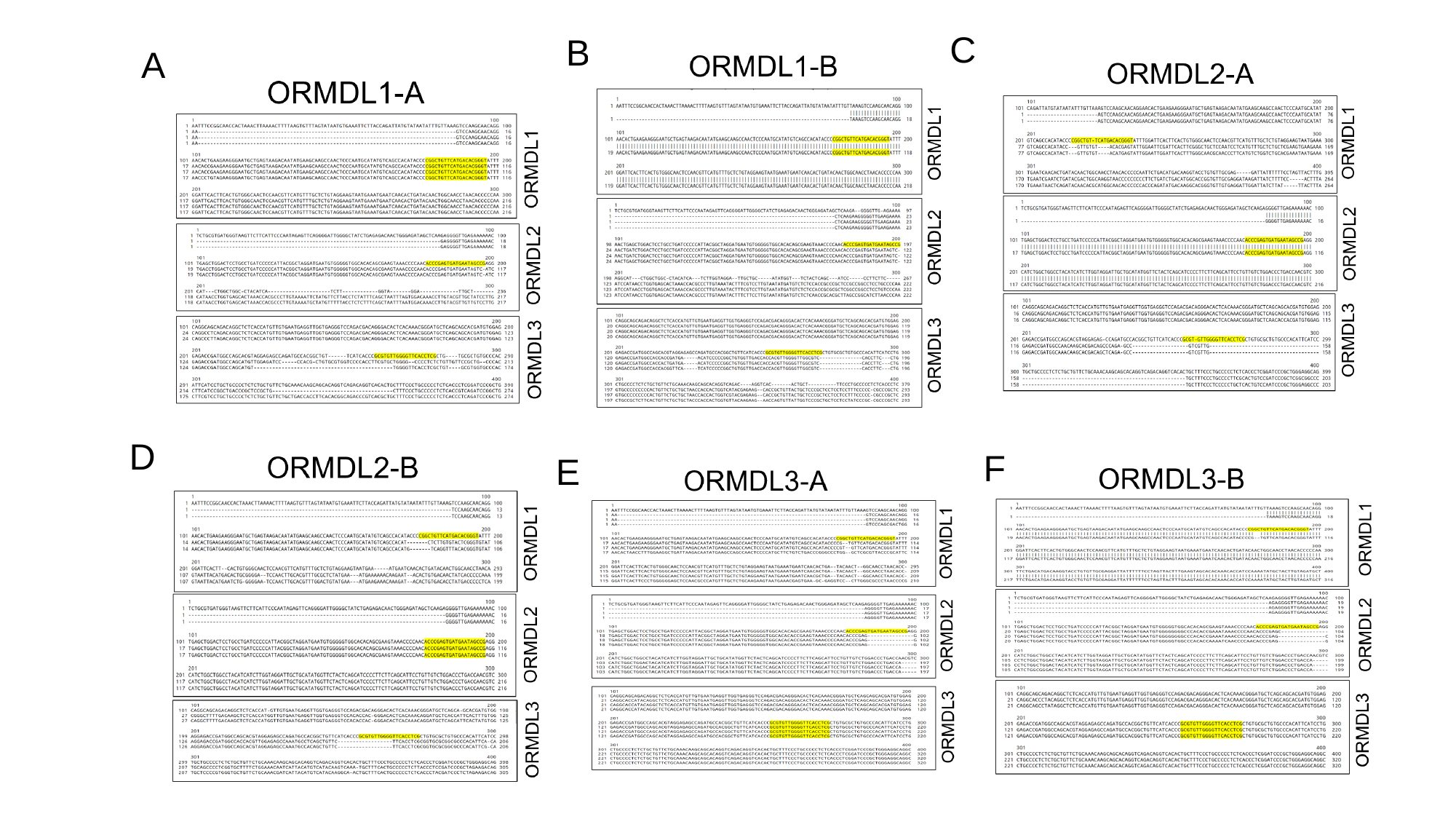

C
B
A
D
F
E

### Slide 3
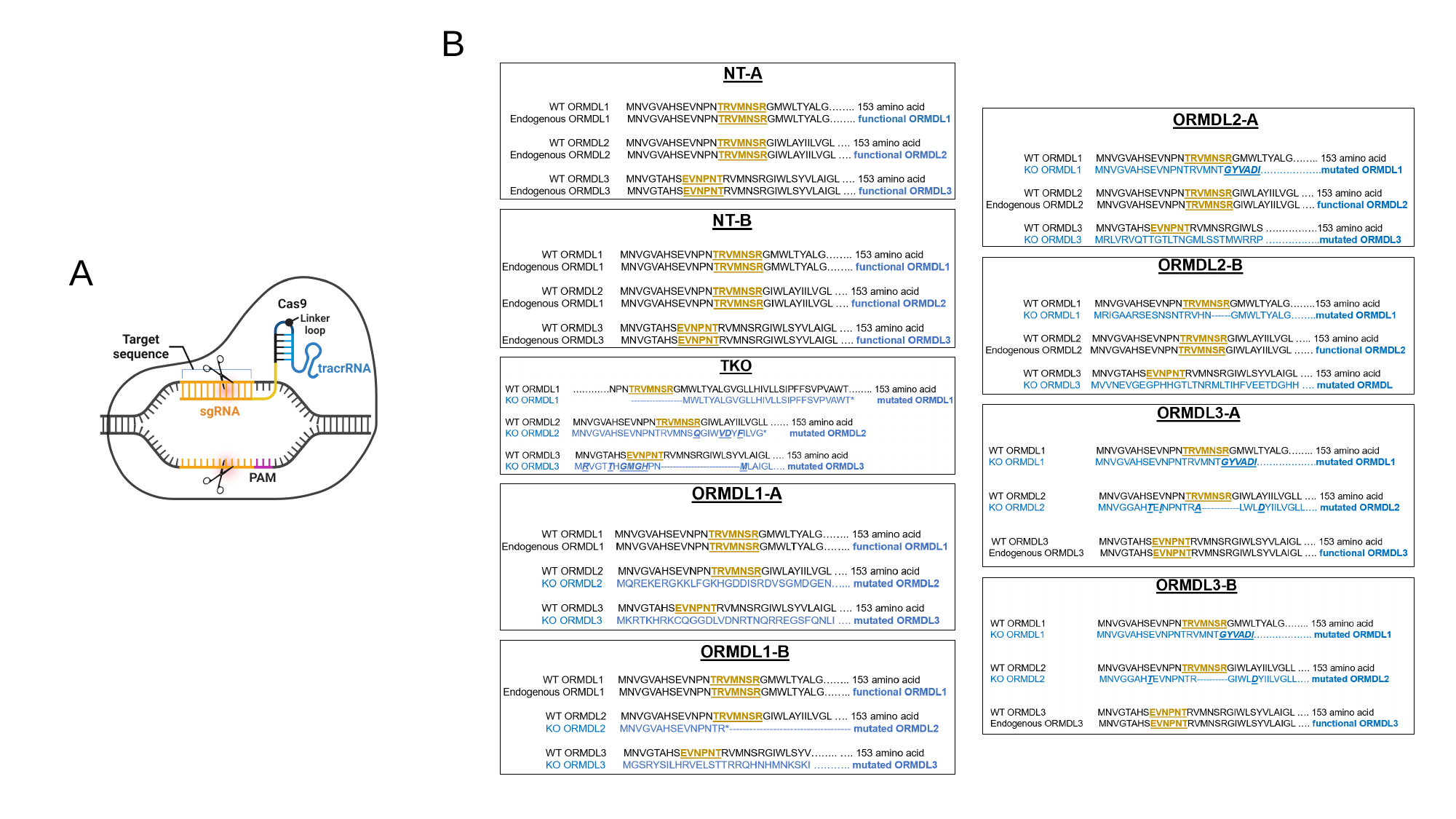

B
A

### Slide 4
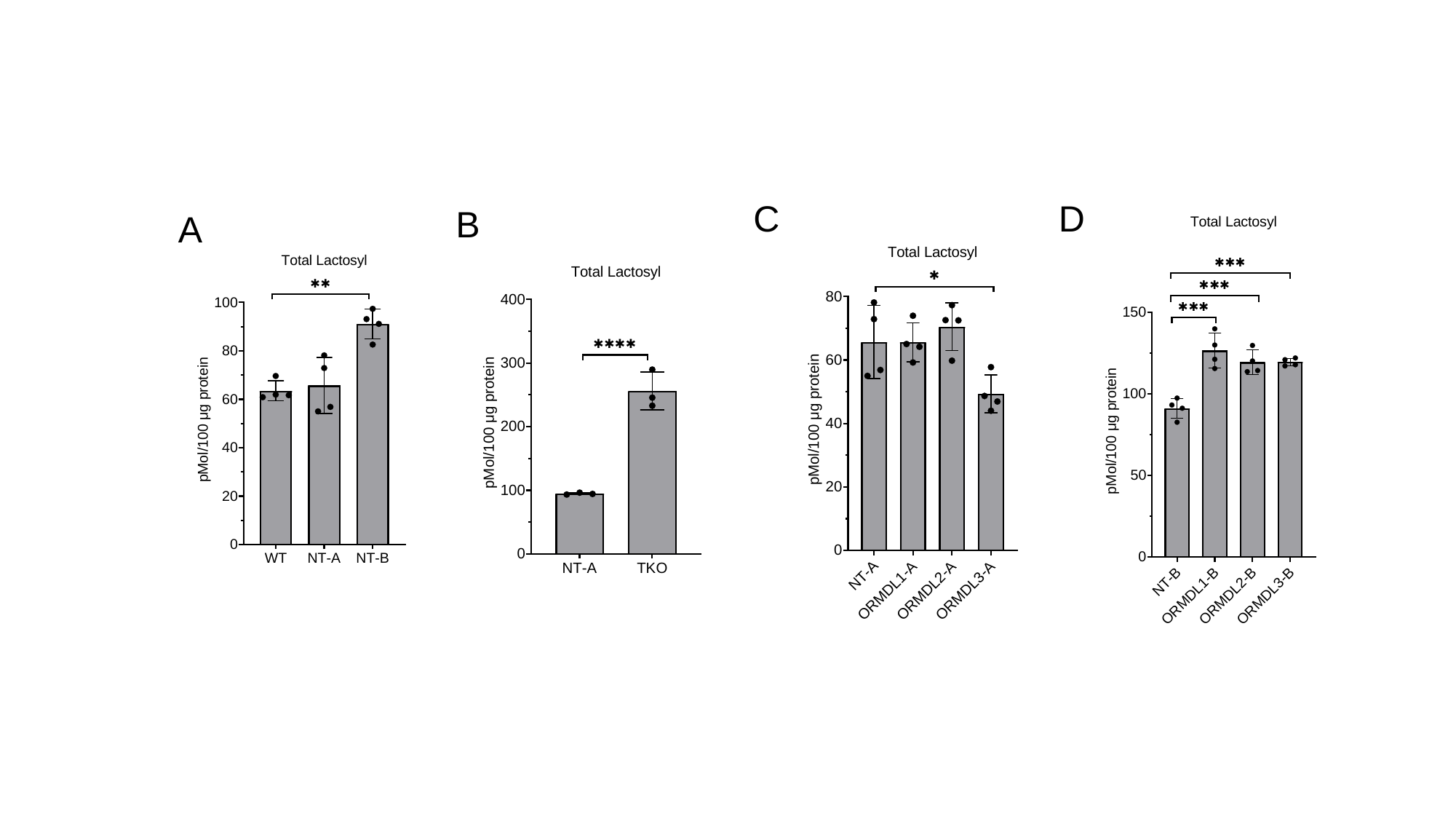

C
D
B
A

### Slide 5
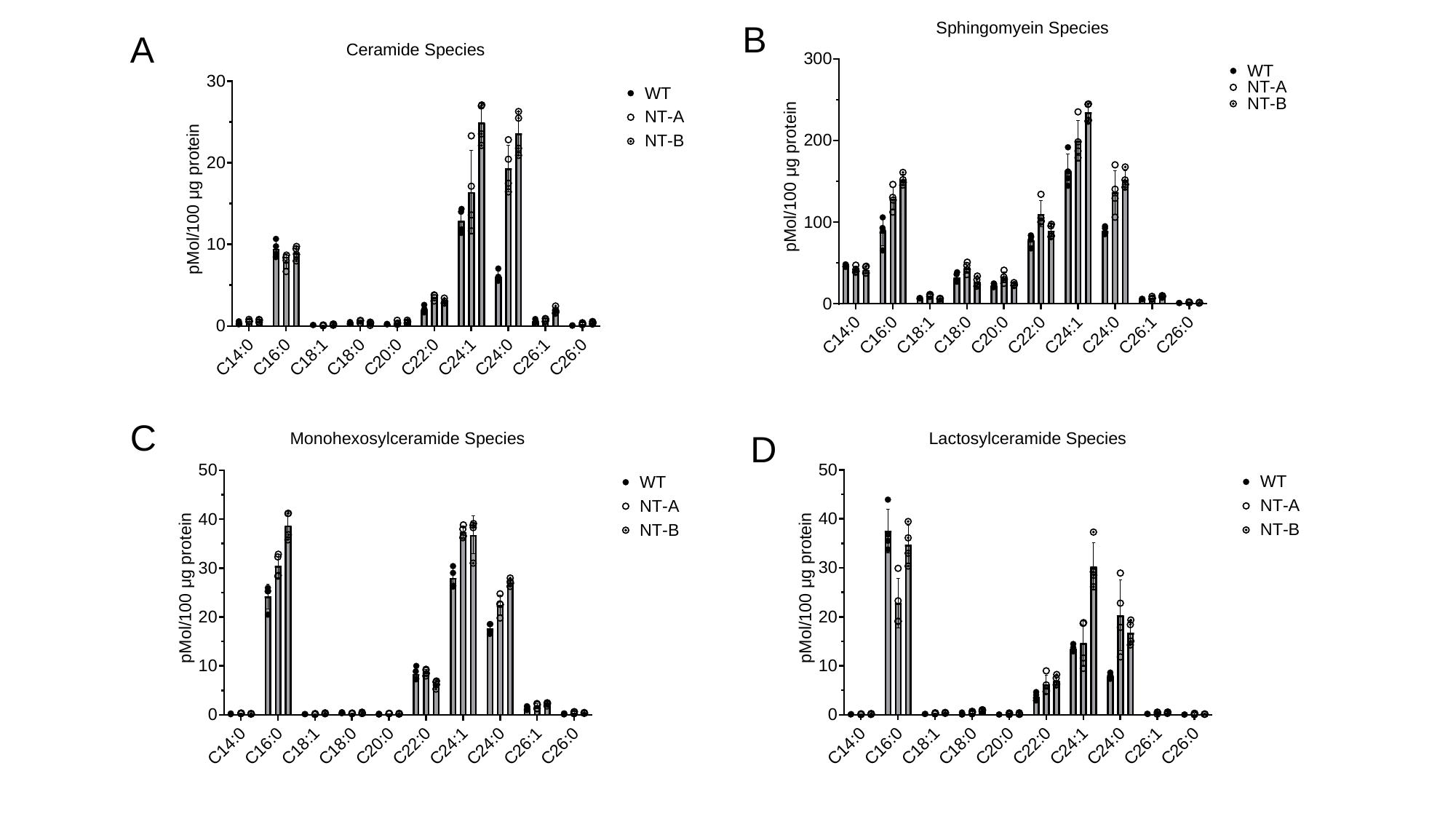

B
A
C
D

### Slide 6
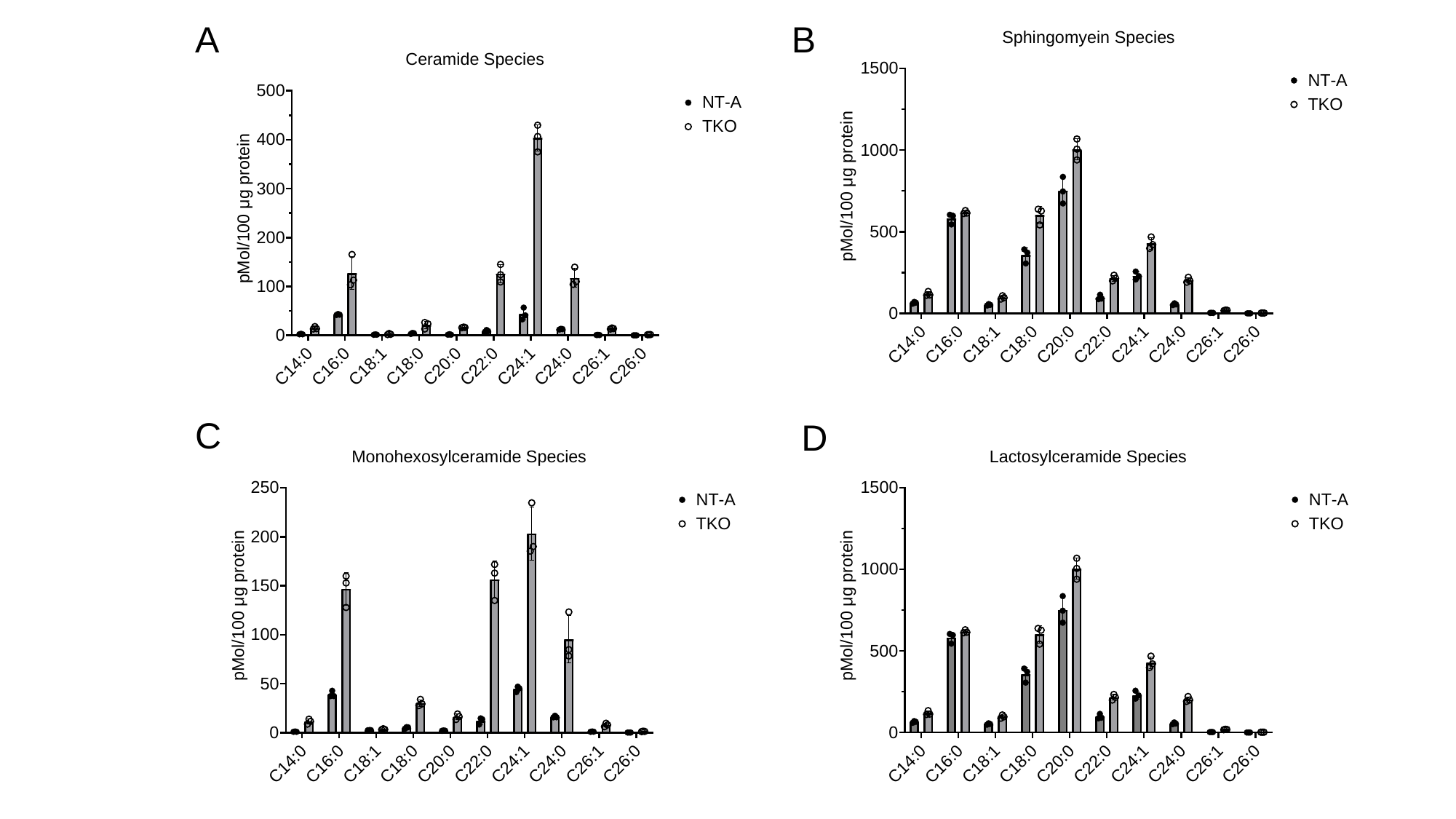

A
B
C
D

### Slide 7
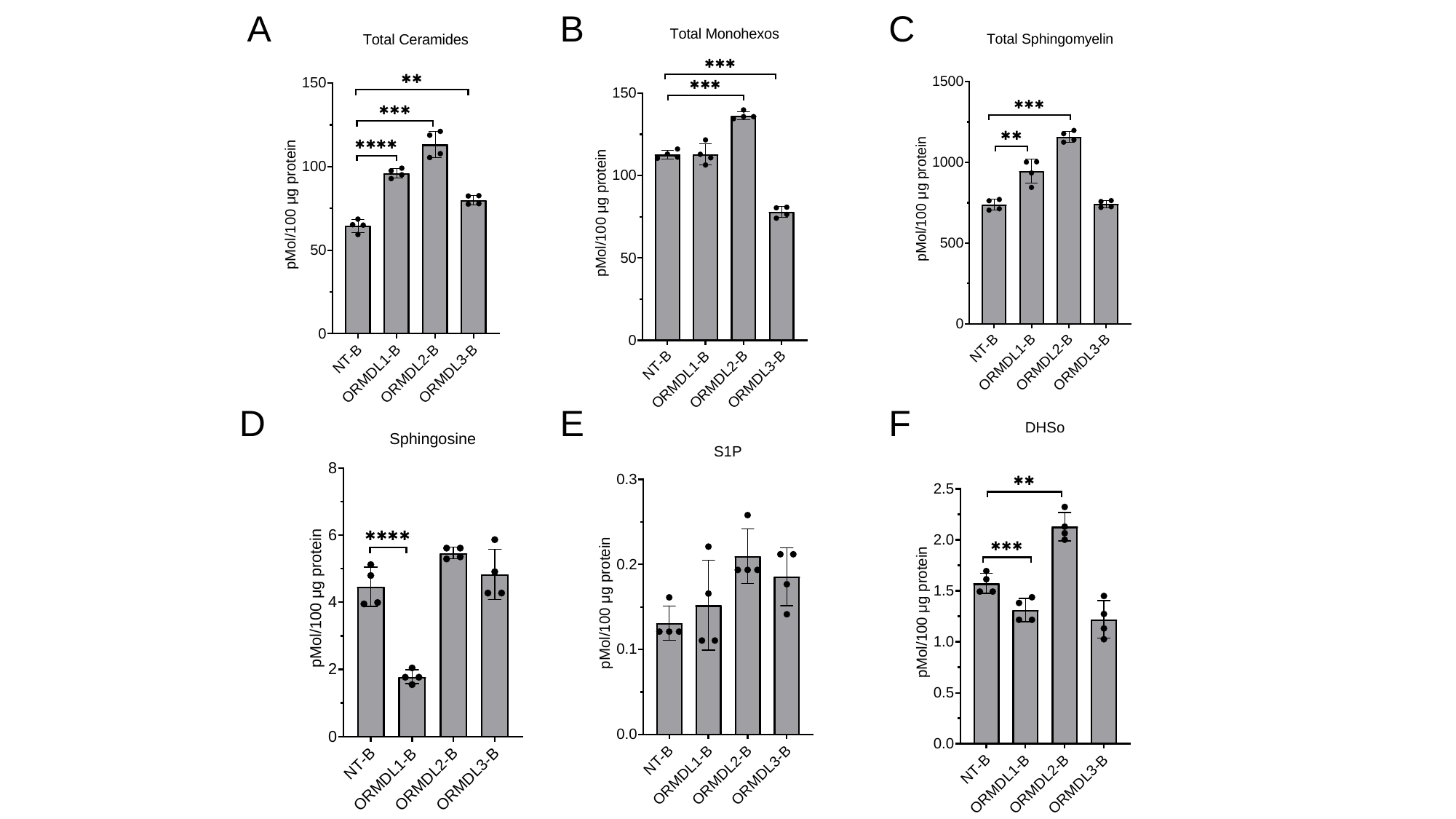

A
B
C
D
E
F

### Slide 8
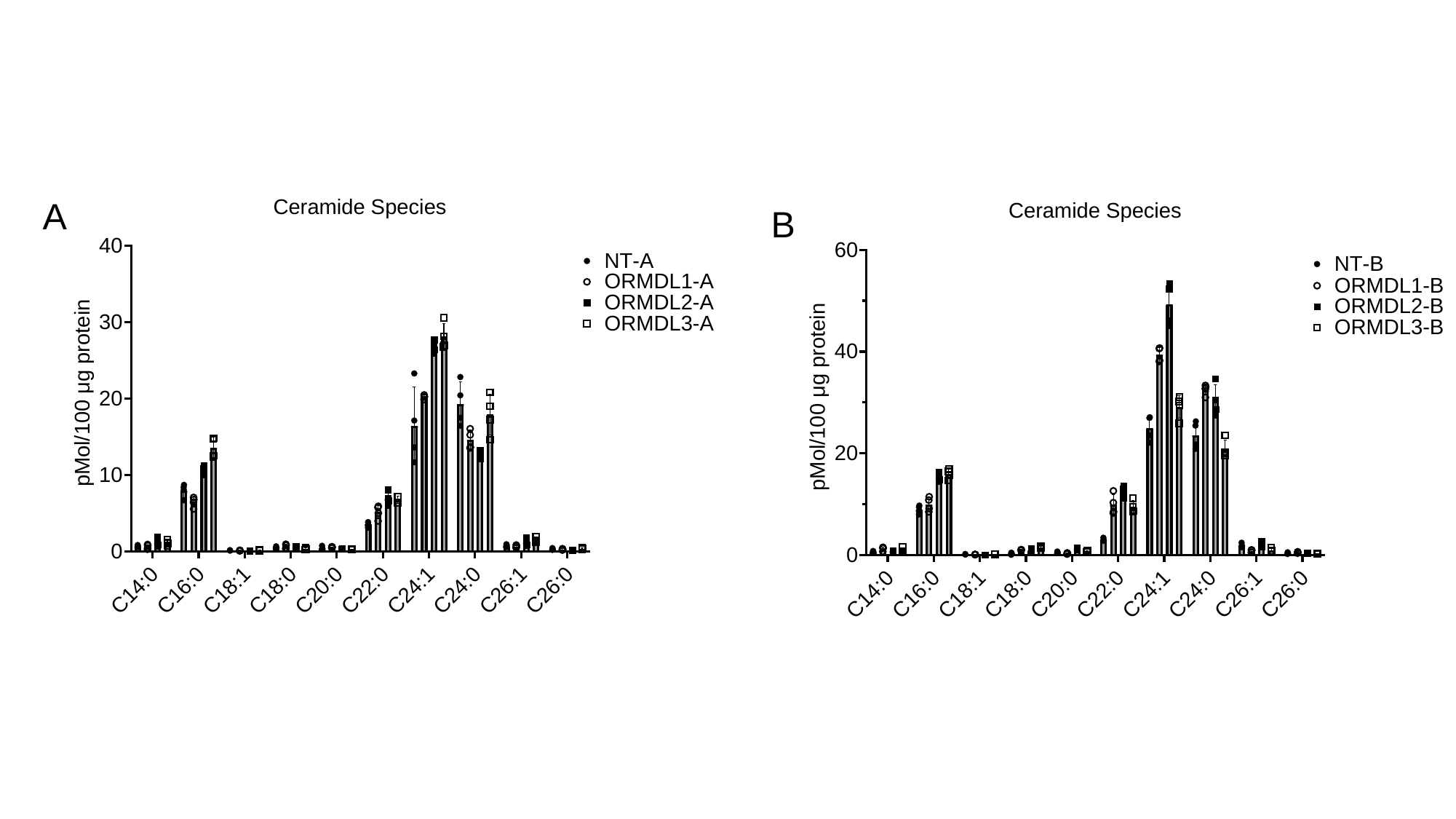

A
B

### Slide 9
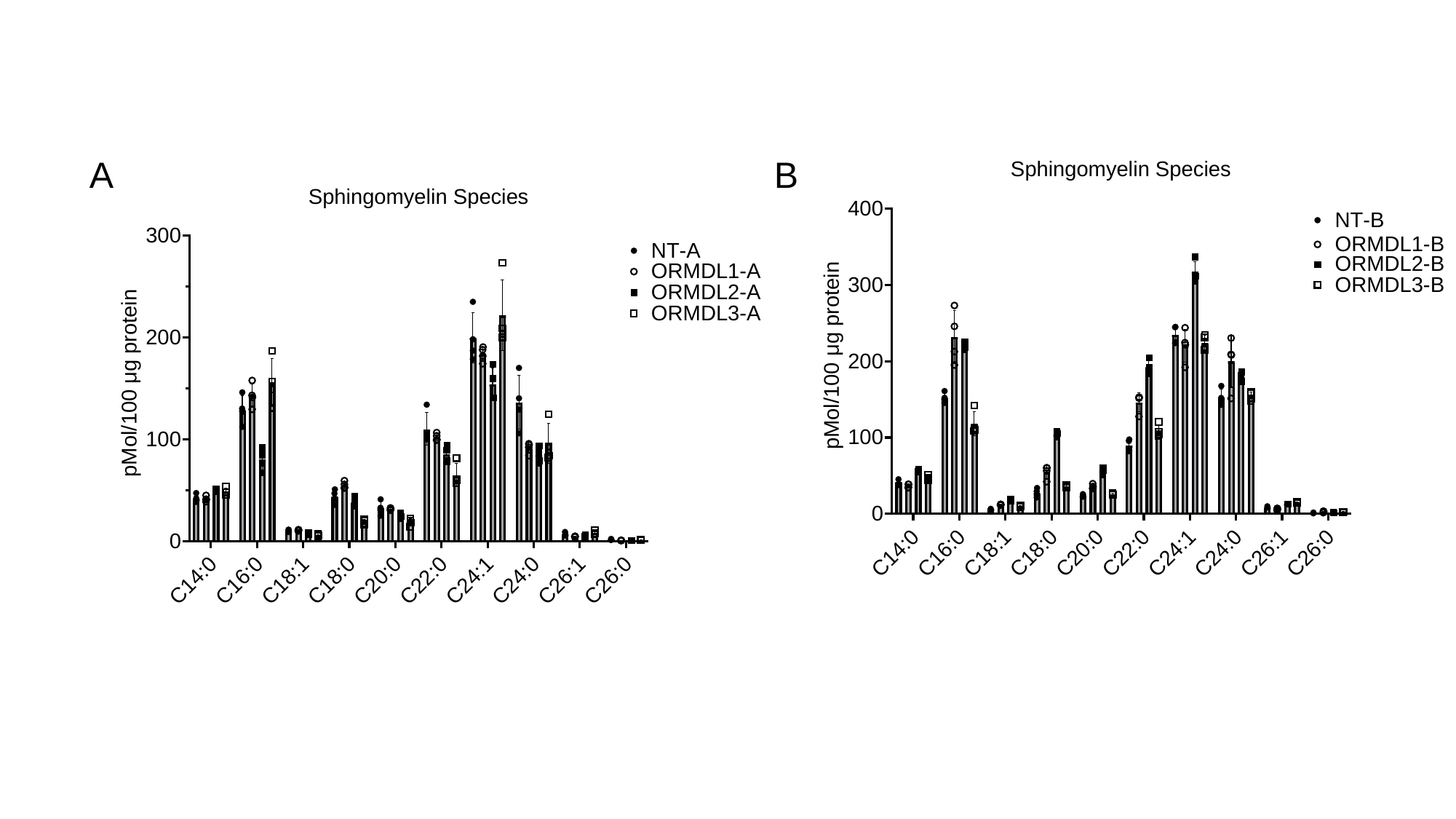

B
A

### Slide 10
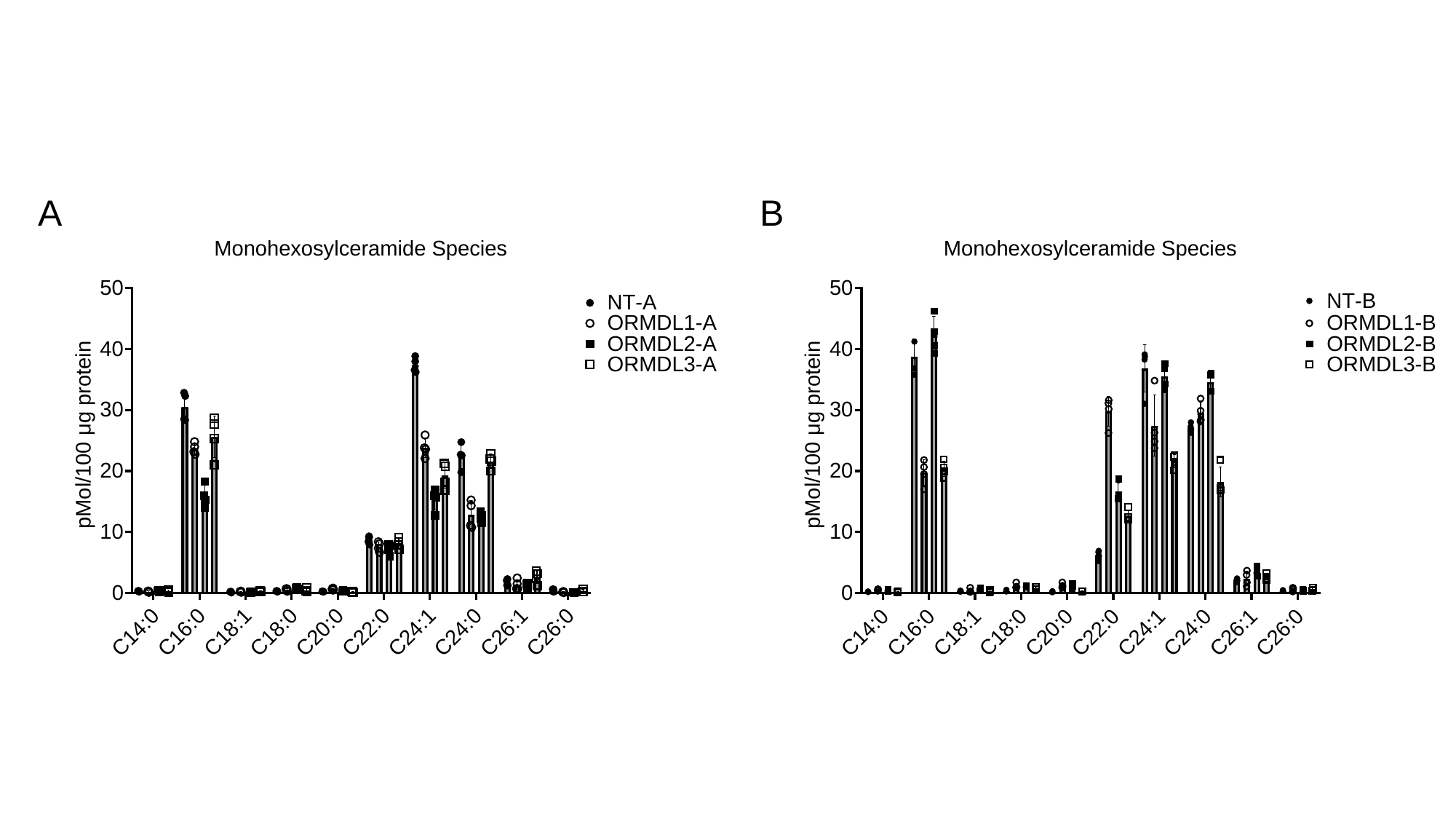

A
B

### Slide 11
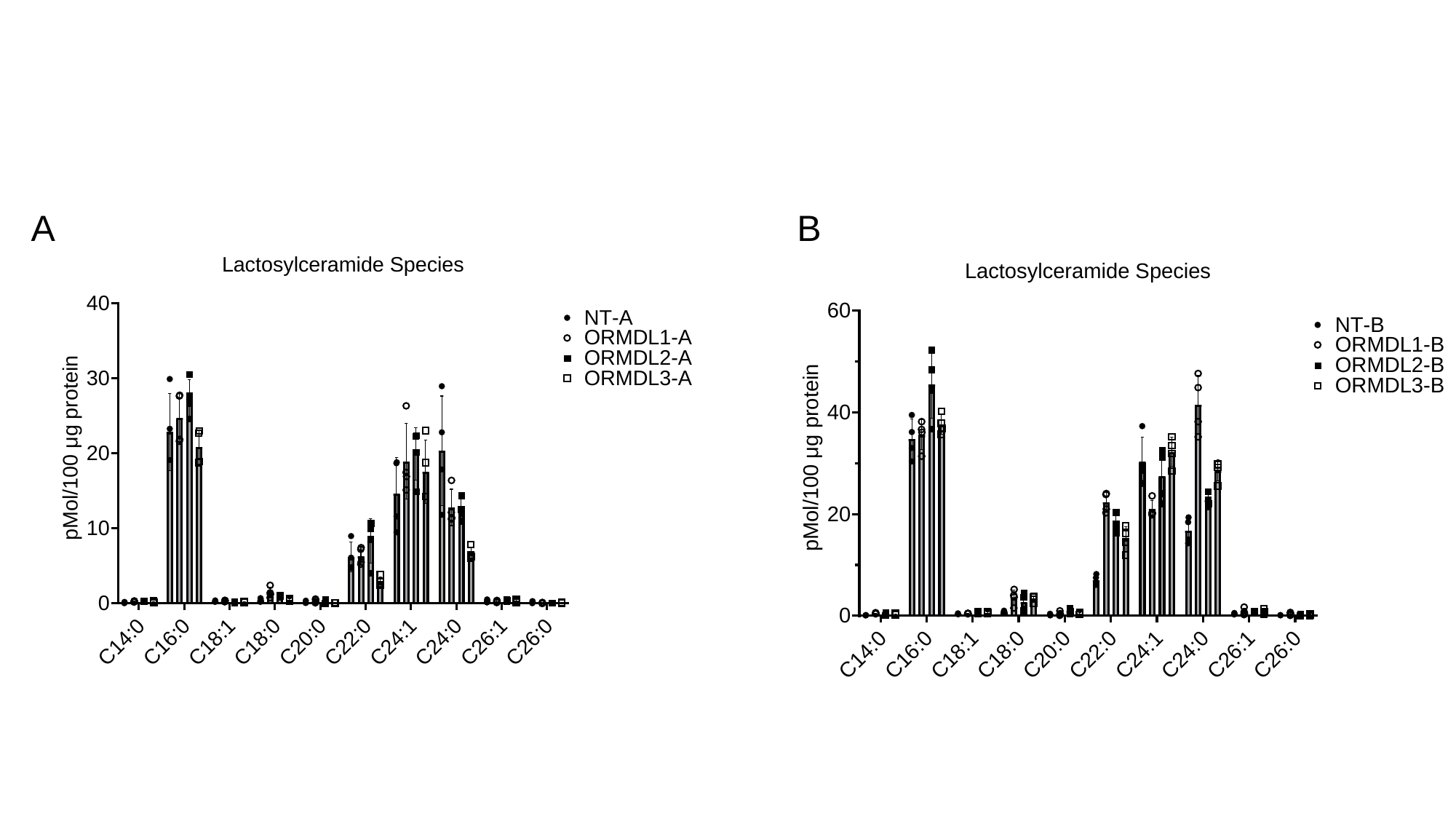

A
B

### Slide 12
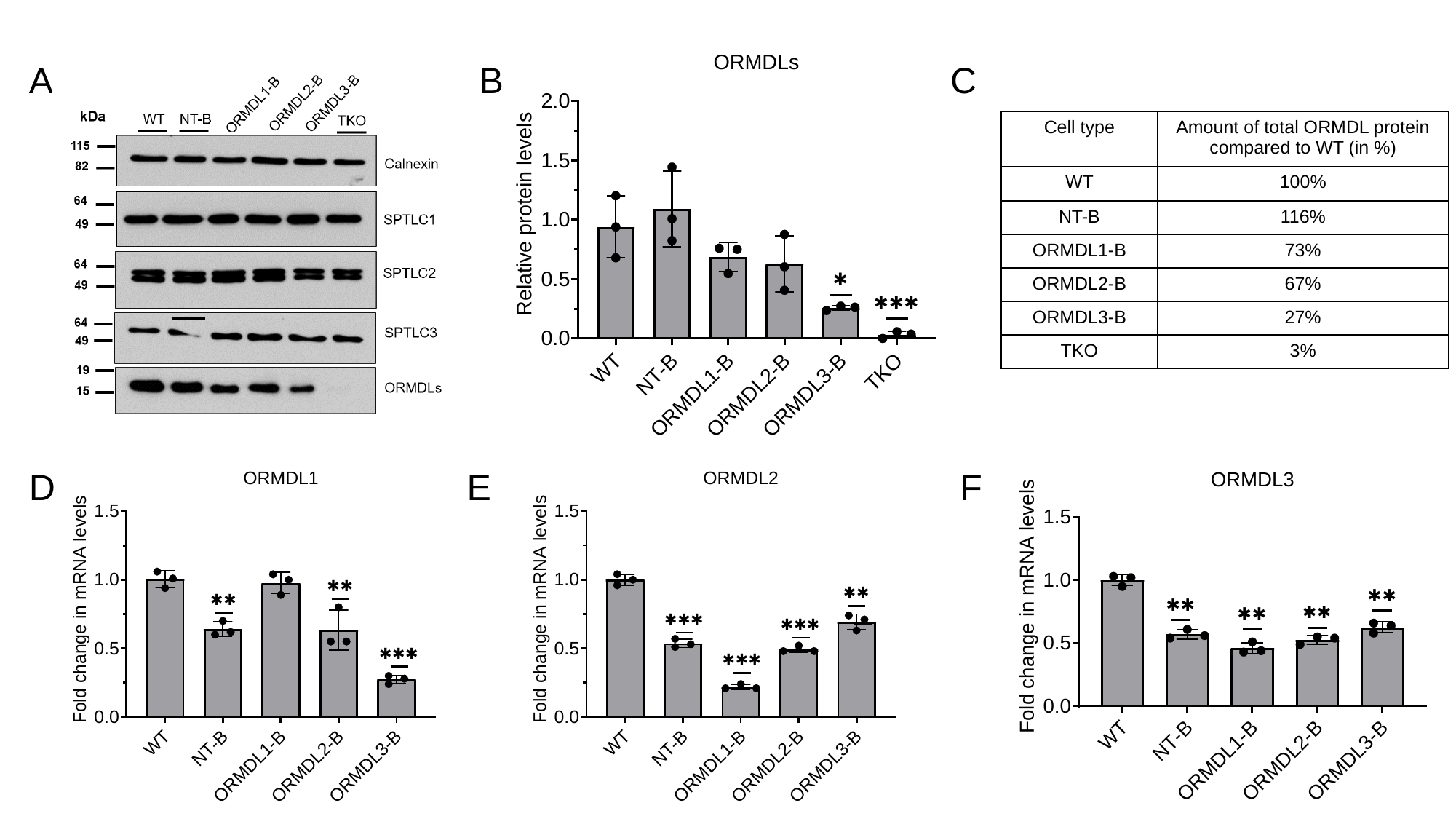

B
C
A
| Cell type | Amount of total ORMDL protein compared to WT (in %) |
| --- | --- |
| WT | 100% |
| NT-B | 116% |
| ORMDL1-B | 73% |
| ORMDL2-B | 67% |
| ORMDL3-B | 27% |
| TKO | 3% |
E
F
D

### Slide 13
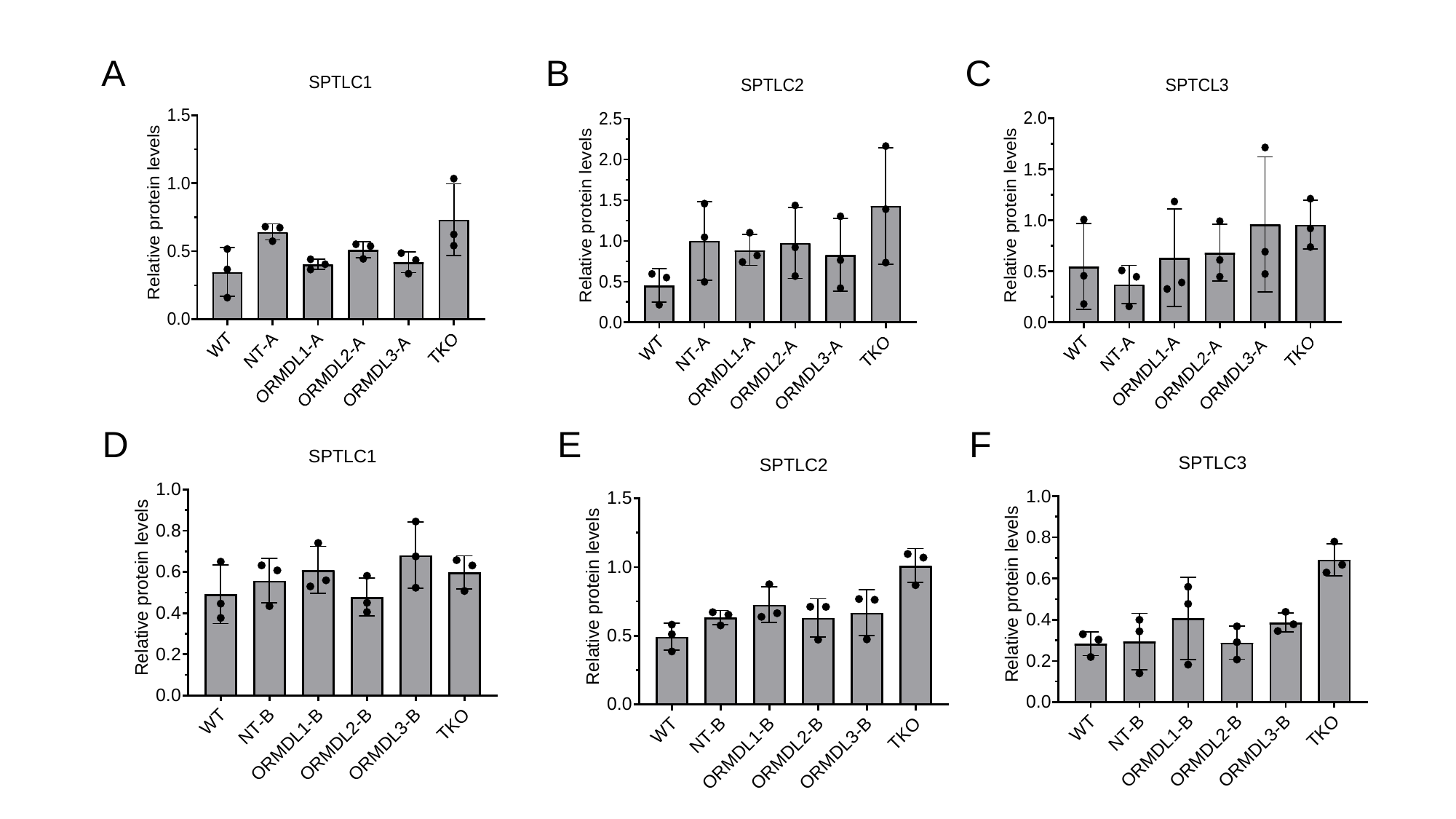

A
B
C
D
E
F

### Slide 14
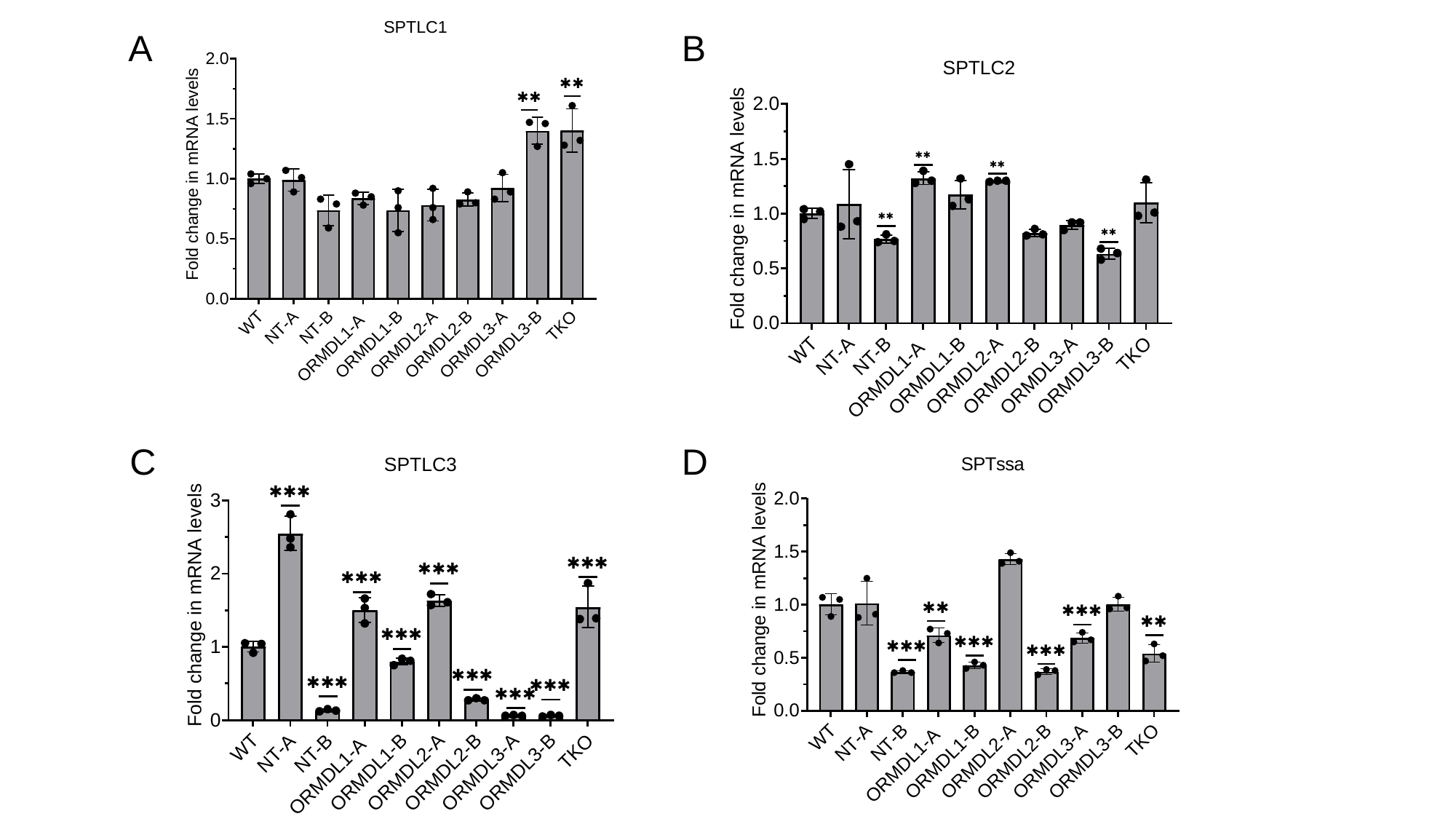

A
B
C
D

### Slide 15
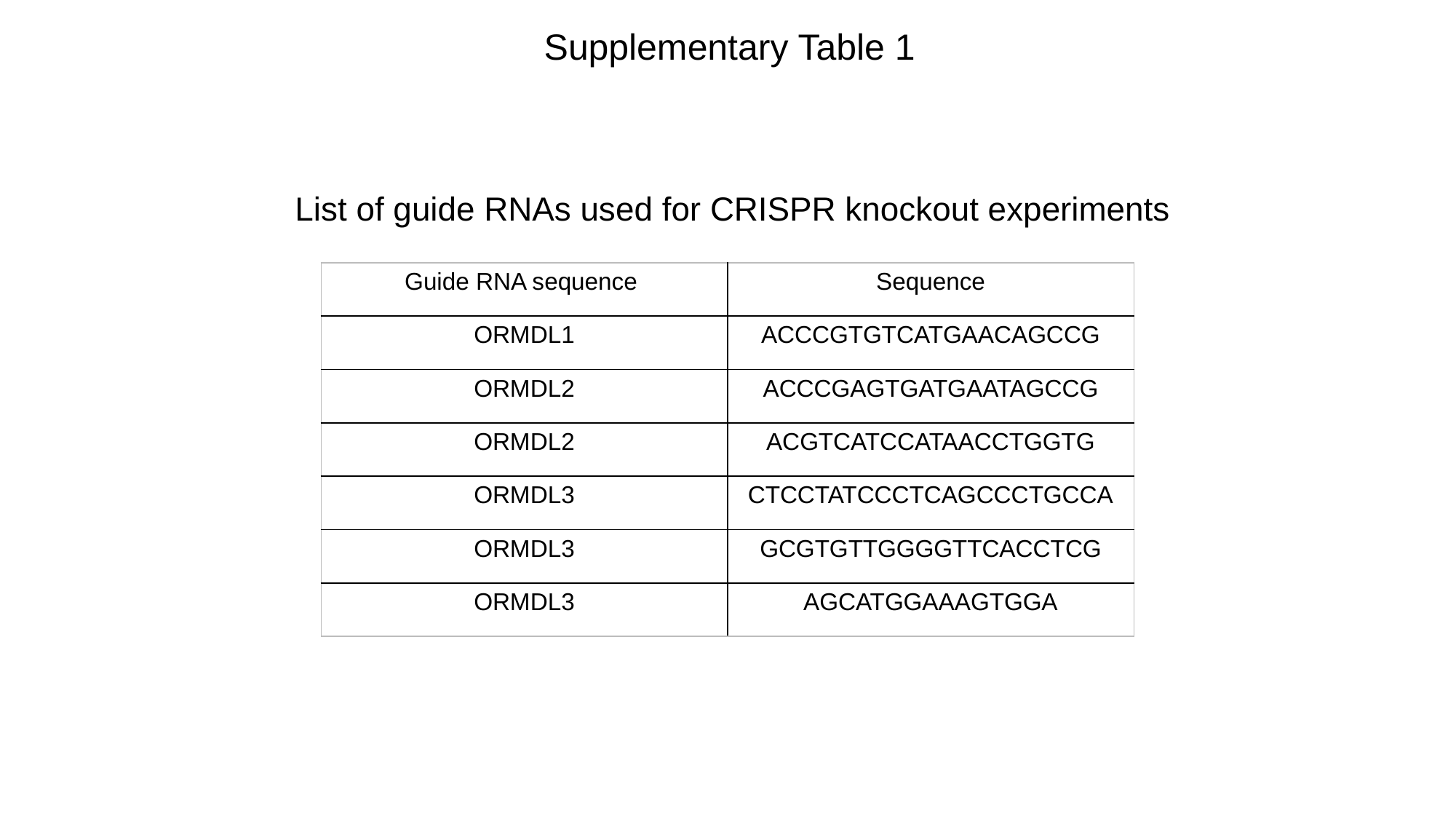

Supplementary Table 1
List of guide RNAs used for CRISPR knockout experiments
| Guide RNA sequence | Sequence |
| --- | --- |
| ORMDL1 | ACCCGTGTCATGAACAGCCG |
| ORMDL2 | ACCCGAGTGATGAATAGCCG |
| ORMDL2 | ACGTCATCCATAACCTGGTG |
| ORMDL3 | CTCCTATCCCTCAGCCCTGCCA |
| ORMDL3 | GCGTGTTGGGGTTCACCTCG |
| ORMDL3 | AGCATGGAAAGTGGA |

### Slide 16
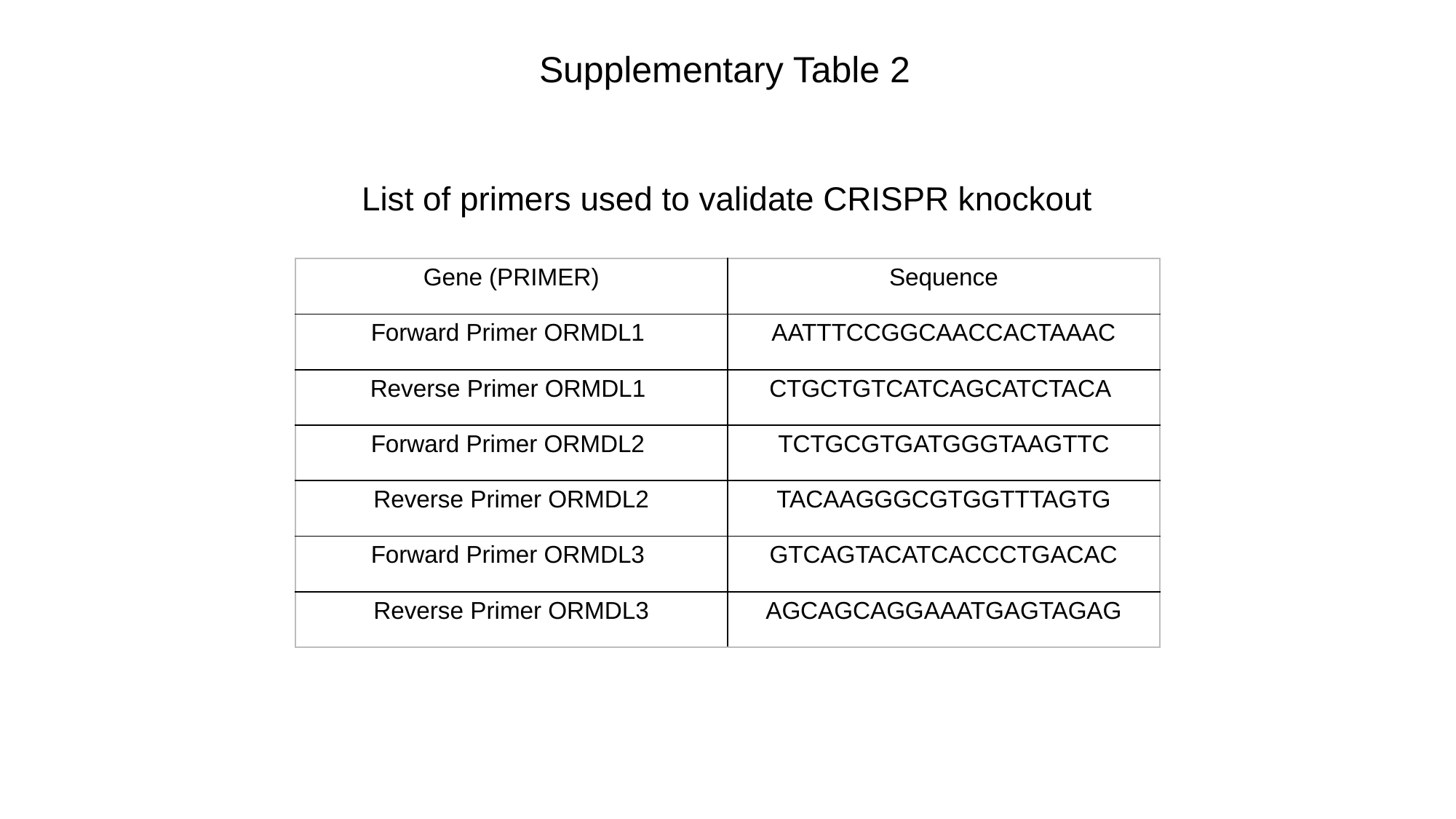

Supplementary Table 2
List of primers used to validate CRISPR knockout
| Gene (PRIMER) | Sequence |
| --- | --- |
| Forward Primer ORMDL1 | AATTTCCGGCAACCACTAAAC |
| Reverse Primer ORMDL1 | CTGCTGTCATCAGCATCTACA |
| Forward Primer ORMDL2 | TCTGCGTGATGGGTAAGTTC |
| Reverse Primer ORMDL2 | TACAAGGGCGTGGTTTAGTG |
| Forward Primer ORMDL3 | GTCAGTACATCACCCTGACAC |
| Reverse Primer ORMDL3 | AGCAGCAGGAAATGAGTAGAG |

### Slide 17
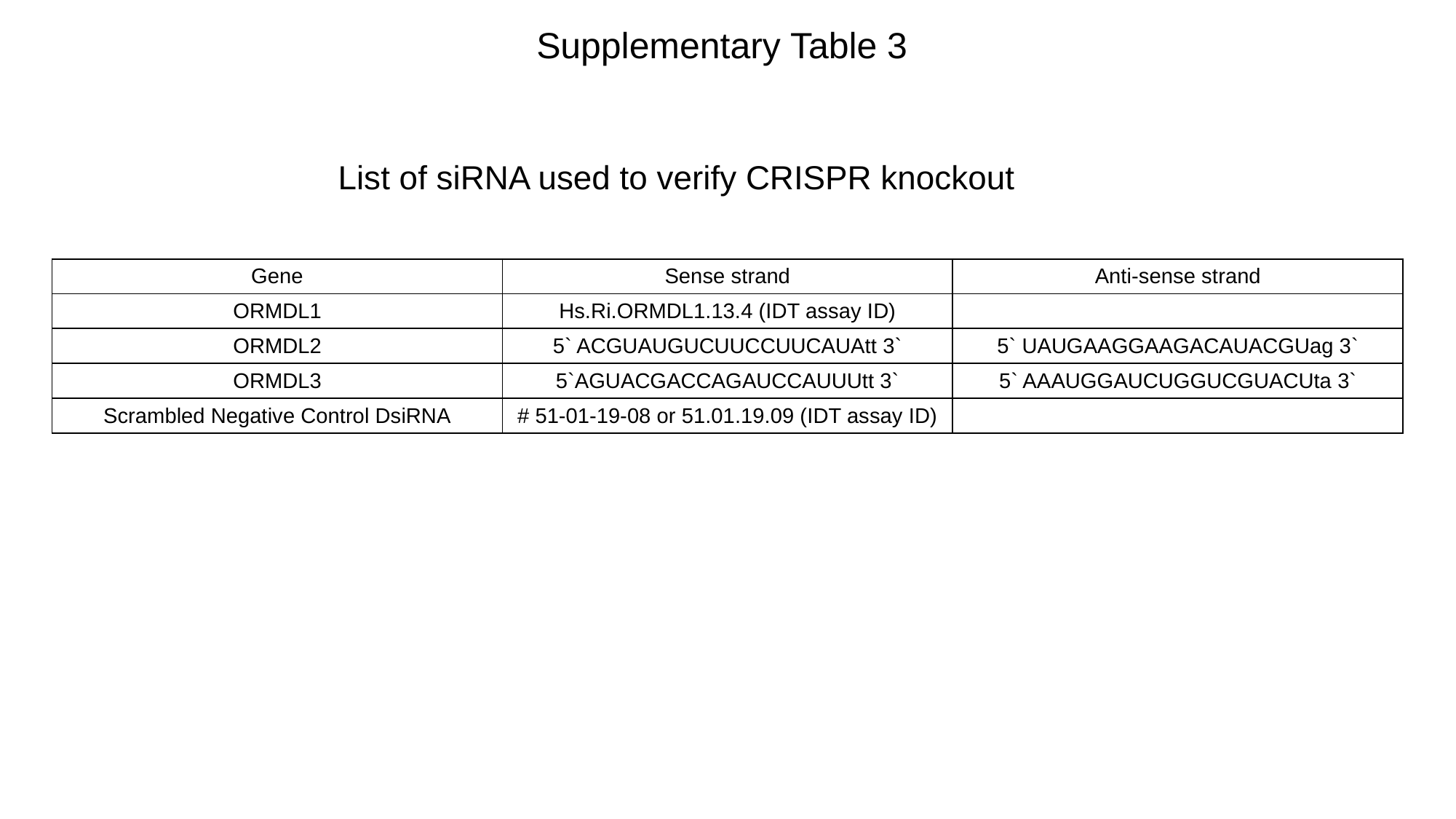

Supplementary Table 3
List of siRNA used to verify CRISPR knockout
| Gene | Sense strand | Anti-sense strand |
| --- | --- | --- |
| ORMDL1 | Hs.Ri.ORMDL1.13.4 (IDT assay ID) | |
| ORMDL2 | 5` ACGUAUGUCUUCCUUCAUAtt 3` | 5` UAUGAAGGAAGACAUACGUag 3` |
| ORMDL3 | 5`AGUACGACCAGAUCCAUUUtt 3` | 5` AAAUGGAUCUGGUCGUACUta 3` |
| Scrambled Negative Control DsiRNA | # 51-01-19-08 or 51.01.19.09 (IDT assay ID) | |

### Slide 18
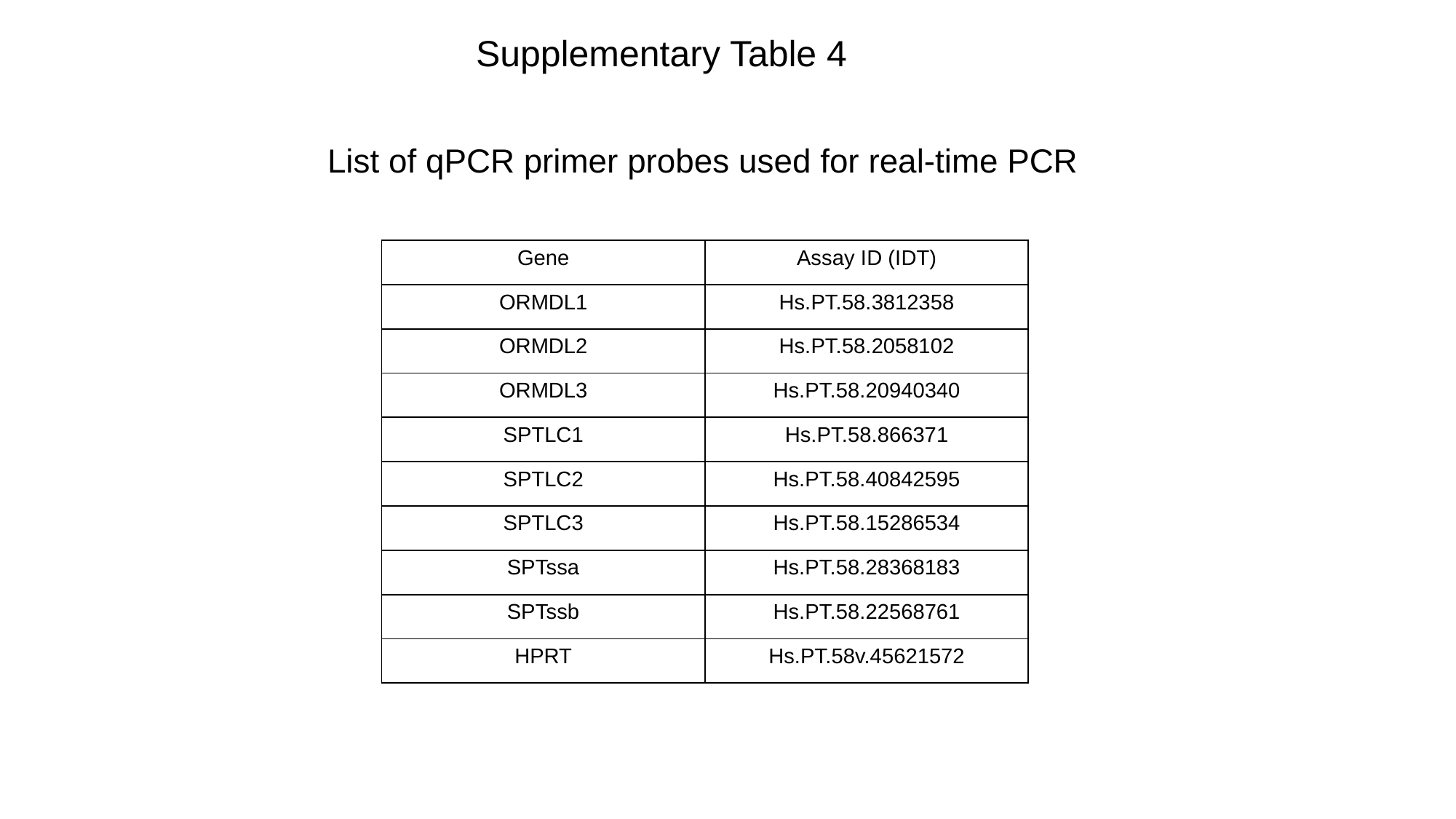

Supplementary Table 4
List of qPCR primer probes used for real-time PCR
| Gene | Assay ID (IDT) |
| --- | --- |
| ORMDL1 | Hs.PT.58.3812358 |
| ORMDL2 | Hs.PT.58.2058102 |
| ORMDL3 | Hs.PT.58.20940340 |
| SPTLC1 | Hs.PT.58.866371 |
| SPTLC2 | Hs.PT.58.40842595 |
| SPTLC3 | Hs.PT.58.15286534 |
| SPTssa | Hs.PT.58.28368183 |
| SPTssb | Hs.PT.58.22568761 |
| HPRT | Hs.PT.58v.45621572 |
